# Cohesin is required for meiotic spindle assembly independent of its role in cohesion in C. elegans

**DOI:** 10.1101/2022.03.10.483728

**Authors:** Karen Perry McNally, Brennan Danlasky, Consuelo Barroso, Ting Gong, Wenzhe Li, Enrique Martinez-Perez, Francis J. McNally

## Abstract

Accurate chromosome segregation requires a cohesin-mediated physical attachment between chromosomes that are to be segregated apart, and a bipolar spindle with microtubule plus ends emanating from exactly two poles toward the paired chromosomes. We asked whether the striking bipolar structure of *C. elegans* meiotic chromosomes is required for bipolarity of acentriolar female meiotic spindles by analyzing mutants that lack cohesion between chromosomes. Both a *spo-11, rec-8, coh-3, coh-4* quadruple mutant and a *spo-11, rec-8* double mutant entered M phase with single chromatids lacking any cohesion. However, the quadruple mutant formed an apolar spindle whereas the double mutant formed a bipolar spindle that segregated chromatids into two roughly equal masses. Residual non-cohesive COH-3/4-dependent cohesin on single chromatids of the double mutant was sufficient to recruit haspin-dependent Aurora B kinase, which regulated the localization of the spindle-assembly factors CLASP-2 and kinesin-13 to mediate bipolar spindle assembly in the apparent absence of chromosomal bipolarity. These results demonstrate that cohesin is essential for spindle assembly and chromosome segregation independent of its role in sister chromatid cohesion.

## Introduction

The accurate segregation of chromosomes during meiosis and mitosis requires sister chromatid cohesion (SCC) provided by the cohesin complex and a bipolar spindle with microtubule minus ends oriented toward the two poles and microtubule plus ends extending from the two poles toward the chromosomes (Nasmyth, 2002). During mitosis in most animal cells, spindle formation is initiated when organelles known as centrosomes are duplicated and move to opposite sides of the cell. There they anchor, nucleate and stabilize microtubules with their plus ends polymerizing away from the poles (Blanco-Ameijeiras, Lozano-Fernandez, & Marti, 2022). Microtubule plus ends puncture the nuclear membrane and capture the kinetochores of chromosomes, thus establishing a symmetric spindle axis.

In contrast to the pathway of mitotic spindle formation, the female meiotic cells of many animals lack centrosomes and spindle formation initiates when microtubules organize around chromatin during the two consecutive meiotic divisions. In *Xenopus* egg extracts and mouse oocytes, DNA-coated beads are sufficient to induce bipolar spindle assembly (Deng, Suraneni, Schultz, & Li, 2007; Heald et al., 1996). The mechanisms of acentrosomal spindle assembly are being elucidated in several species and two alternate pathways have been implicated. The first molecular activity to be identified in the assembly of microtubules around meiotic chromatin is the GTPase Ran. In the Ran pathway, spindle assembly factors (SAFs) contain nuclear localization sequences and are imported into the nucleus during interphase by binding to importins. GTP-ran, which is maintained at a high concentration in the nucleus by the chromatin-bound GEF RCC1, causes dissociation of the SAFs from importins inside the nucleus, thus driving the directionality of import. Upon nuclear envelope breakdown, tubulin enters the region adjacent to chromatin and the locally activated SAFs initiate MT nucleation and stabilization (Cavazza & Vernos, 2015). Inhibition of the Ran pathway prevents or affects the assembly of acentrosomal spindles in *Xenopus* egg extracts (Carazo-Salas et al., 1999) and in mouse (Dumont et al., 2007), *Drosophila* (Cesario & McKim, 2011) and *C. elegans* oocytes (Chuang, Schlientz, Yang, & Bowerman, 2020). In *Xenopus* egg extracts, spindle assembly is induced by beads coated with the ran GEF, RCC1, even without DNA (Halpin, Kalab, Wang, Weis, & Heald, 2011).

The second pathway which has been implicated in acentrosomal spindle assembly requires the Chromosomal Passenger Complex (CPC), which includes the chromatin-targeting proteins Survivin and Borealin, the scaffold subunit INCENP, and Aurora B kinase (Willems et al., 2018). The CPC is recruited to distinct regions on mitotic chromosomes by at least three different pathways (Broad, DeLuca, & DeLuca, 2020). Depletion of CPC components resulted in a lack of spindle microtubules in *Drosophila* oocytes (Radford, Jang, & McKim, 2012) and in *Xenopus* egg extracts to which sperm nuclei or DNA-coated beads are added (Kelly et al., 2007; Maresca et al., 2009; Sampath et al., 2004). In *C. elegans* oocytes, the CPC subunits, BIR-1/survivin (Speliotes, Uren, Vaux, & Horvitz, 2000), INCENP (Wignall & Villeneuve, 2009), and the Aurora B-homolog AIR-2 (Divekar, Davis-Roca, Zhang, Dernburg, & Wignall, 2021; Dumont, Oegema, & Desai, 2010) contribute to meiotic spindle assembly.

While the GTP Ran and CPC pathways are known to be involved in the initiation of acentrosomal spindle assembly, the mechanism by which the microtubules are captured into two poles is unclear. Spindles with one or more poles form when chromatin-coated beads are added to *Xenopus* egg extracts, suggesting that pole formation is an intrinsic activity of microtubules assembling around chromatin (Halpin et al., 2011). However, the results also suggest that the reproducible production of bipolar spindles requires that the process includes some bidirectionality. In *C. elegans*, meiotic bivalents, which promote assembly of a bipolar metaphase I spindle, are composed of 4 chromatids held together by chiasmata, physical attachments provided by cohesin and a single crossover formed between homologous chromosomes. These bivalents have a discrete bipolar symmetry with a mid-bivalent ring containing the CPC, and they are capped at their two ends by cup-shaped kinetochores. Metaphase II univalents, which promote assembly of a bipolar metaphase II spindle, are composed of 2 chromatids held together by cohesin. These univalents also have a discrete bipolar symmetry with a CPC ring between sister chromatids that are each capped by cup-shaped kinetochores (Dumont et al., 2010; Monen, Maddox, Hyndman, Oegema, & Desai, 2005; Wignall & Villeneuve, 2009).

To test whether this chromosomal bipolar symmetry is required for spindle bipolarity, we analyzed cohesin mutants that start meiotic spindle assembly with single chromatids rather than the bivalents present in wild-type meiosis I or the univalents present in wild-type meiosis II. During meiosis, cohesin is composed of SMC-1, SMC-3, and one of 3 meiosis-specific kleisin subunits: REC-8 and the highly identical and functionally redundant COH-3 and COH-4 (Pasierbek et al., 2001; Severson, Ling, van

Zuylen, & Meyer, 2009; Severson & Meyer, 2014). Both REC-8 and COH-3/4 cohesin promote pairing and recombination between homologous chromosomes during early meiosis, thus ensuring chiasma formation. However, SCC appears to be provided by REC-8 complexes, while COH-3/4 complexes associate with indivual chromatids (Crawley et al., 2016; Woglar et al., 2020). Previous work indicated that *rec-8* single mutants have 12 univalents at meiosis I, with each pair of sister chromatids held together by recombination events dependent on COH-3/COH-4 cohesin (Cahoon, Helm, & Libuda, 2019; Crawley et al., 2016). Sister chromatids segregated equationally at anaphase I of *rec-8* mutants with half the chromatids going into a single polar body (Severson et al., 2009). This suggests that *rec-8* embryos enter metaphase II with 12 single chromatids. Although it was reported that *rec-8* embryos do not extrude a second polar body, the structure of the metaphase II spindle was not described in detail. To address the question of whether chromosomal bipolarity is required for spindle bipolarity, we first monitored metaphase II spindle assembly in a *rec-8* mutant by time-lapse imaging of living embryos *in utero*.

## Results

### Apolar spindles assemble around single chromatids of metaphase II *rec-8* embryos

Time-lapse *in utero* imaging of control embryos with microtubules labelled with mNeonGreen::tubulin and chromosomes labelled with mCherry::histone H2b revealed bipolar spindles that shorten, then rotate, then segregate chromosomes in both meiosis I and meiosis II (Fig. 1A; Video S1). Wild-type embryos enter metaphase I with 6 bivalents and enter metaphase II with 6 univalents whereas *rec-8* embryos enter metaphase I with 12 univalents and enter metaphase II with approximately 12 single chromatids (Fig. 1B) (Severson et al., 2009). Time-lapse imaging of *rec-8* embryos revealed bipolar metaphase I spindles that shortened, rotated, and segregated chromosomes (Fig. 1C, -1:45-5:15; Video S2). Metaphase II *rec-8* embryos, however, assembled an amorphous cloud of microtubules around single chromatids which did not segregate into two masses. The apolar spindle shrank with timing similar to spindle shortening that occurs during wild-type meiosis (Fig. 1C, 9:15-18:00). Because spindle shortening is caused by APC-dependent inactivation of CDK1 (Ellefson & McNally, 2011), this suggests that the failure in metaphase II spindle assembly is not due to a lack of cell cycle progression. The bipolar nature of metaphase I *rec-8* spindles and the apolar nature of *rec-8* metaphase II spindles was confirmed by time-lapse imaging of GFP::ASPM-1 (Fig. 1D). ASPM-1 binds at microtubule minus ends (Jiang et al., 2017) so the dispersed appearance of GFP::ASPM-1 on *rec-8* metaphase II spindles suggests that microtubules are randomly oriented in the spindle.

**Figure 1.**
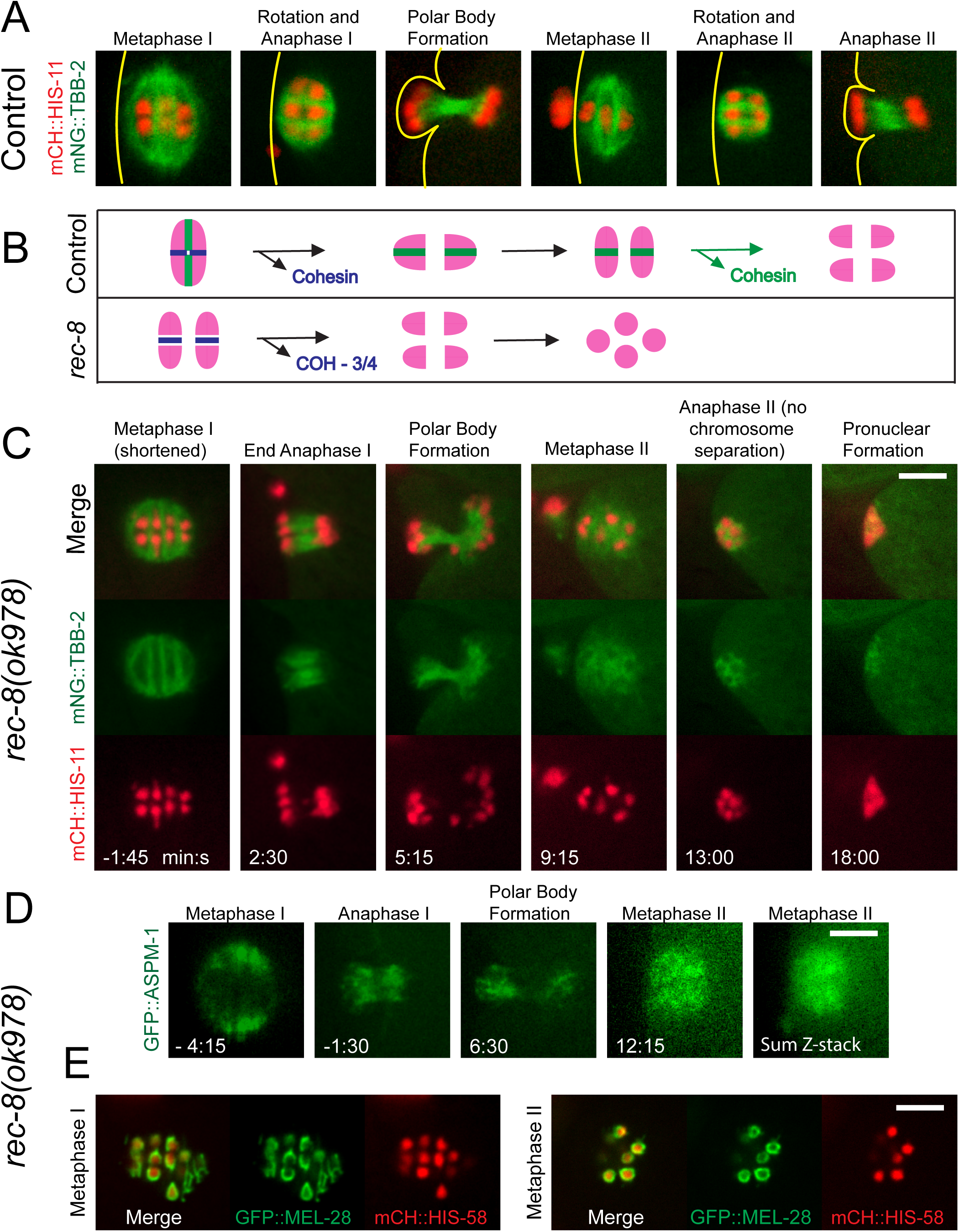
Metaphase II spindles are apolar in *rec-8(ok978)*. **(A)** In control embryos, bipolar MI spindles shorten and rotate, chromosomes segregate, and a polar body forms. The cycle repeats with a bipolar MII spindle. Lines indicate the position of the cortex. **(B)** In Metaphase I, both sister chromatids and homologs are bound by cohesin; homologs are released and separate in Anaphase I; sister chromatids are released and separate in Anaphase II. In *rec-8(ok978)*, COH-3/4 loosely binds sister chromatids in MI and no cohesin is present in MII. **(C)** Time-lapse imaging of *rec-8(ok978)* expressing mNG::TBB-2 and mCH::HIS-11. The metaphase II spindle appears disorganized and no anaphase chromosome separation occurs in 8/8 embryos. 0 minutes is the end of MI spindle rotation. **(D)** Time-lapse imaging of *rec-8(ok978)* expressing GFP::ASPM-1. Single-focal plane imaging was ended at metaphase II and a z-stack acquired. 7/7 metaphase I spindles were bipolar and 8/8 metaphase II spindles were apolar. **(E)** Imaging of *rec-8(ok978)* expressing GFP::MEL-28 revealed kinetochore cups in 4/4 metaphase I spindles and chromatids enclosed by GFP::MEL-28 in 7/7 metaphase II spindles. All bars = 4μm.

Time-lapse imaging of the kinetochore protein GFP::MEL-28 in *rec-8* embryos revealed metaphase I univalents with discrete bipolar structure similar to wild-type metaphase II univalents, whereas metaphase II single chromatids were enveloped by a contiguous symmetrical shell of GFP::MEL-28 (Fig. 1E).

### Apolar spindles assemble around single chromatids of metaphase I *spo-11 rec-8 coh-4 coh-3* embryos

To test whether the apparent inability of single chromatids to drive bipolar spindle assembly is specific for meiosis II, we analyzed embryos of a *spo-11 rec-8 coh-4 coh-3* quadruple mutant (Fig. 2A), which lack meiotic cohesin and the double strand breaks that initiate meiotic recombination (*spo-11* mutation) and therefore enter metaphase I with 24 single chromatids (Severson et al., 2009) (Fig. 2B). In these embryos, an amorphous mass of microtubules formed around the 24 chromatids (Fig. 2A, -2:30; Video S3). This cloud of microtubules shrank with similar timing to wild-type spindle shortening but was not followed by any separation of chromosomes (Fig. 2A, -2:30 -2:30). A second large mass of microtubules formed at the time that a metaphase II spindle normally forms (Fig. 2A, 12:15). This metaphase II mass also shrank with similar timing to normal spindle shortening (Fig. 2A, 12:15-16) but chromatids did not separate into two masses. These results indicated that bipolar spindles cannot assemble around single chromatids that lack both cohesin and cohesion, at both metaphase I and metaphase II.

**Figure 2.**
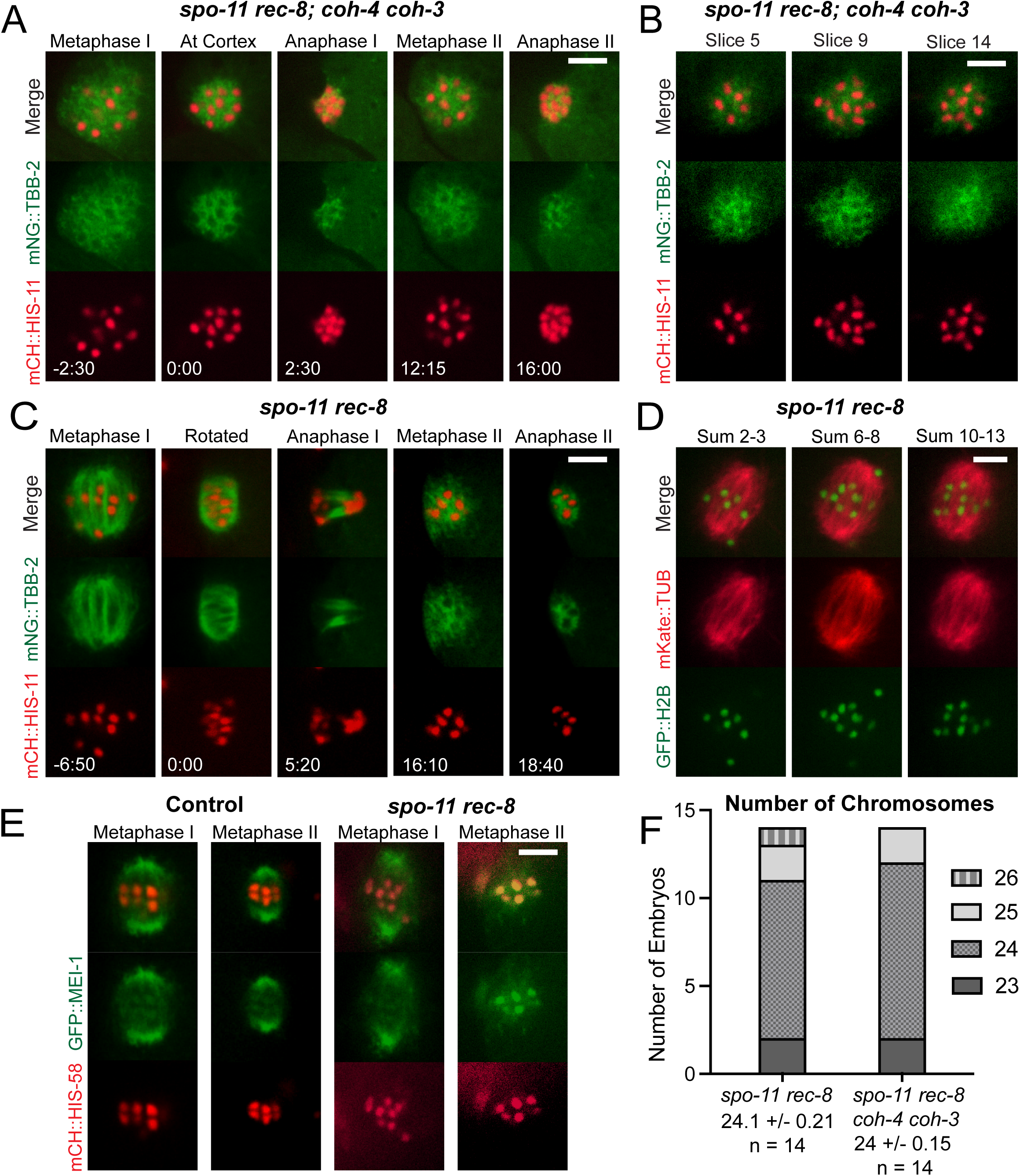
*spo-11 rec-8; coh-4 coh-3* embryos have disorganized meiotic spindles; *spo-11 rec-8 embryos* have bipolar spindles in meiosis I. (A) Single-focal plane time-lapse imaging of a *spo-11 rec-8; coh-4 coh-3* mutant expressing mNeonGreen::TBB-2 and mCherry::HIS. Disorganized spindles were observed in both MI and MII in 10/10 embryos. 0 minutes is the time when the MI spindle contacts the cortex. (B) Z-stack slices of a *spo-11 rec-8; coh-4 coh-3* MI spindle show 24 chromatids with one chromatid visible in both slices 9 and 14. (C) Single-focal plane time-lapse imaging of 13/13 *spo-11 rec-8* embryos show bipolar MI spindles which undergo anaphase chromosome separation and MII spindles which are disorganized and do not undergo anaphase chromosome separation. 0 minutes is the completion of MI spindle rotation. (D) Combined z-stack slices of a *spo-11 rec-8* MI spindle show 24 chromatids. (E) Time-lapse imaging of *spo-11 rec-8* embryos expressing GFP::MEI-1. 10/10 Control MI spindles, 5/5 Control MII spindles and 9/9 *spo-11 rec-8* MI spindles were bipolar. 8/8 *spo-11 rec-8* MII spindles were apolar. (F) Graph showing chromosome numbers during MI in both *spo-11 rec-8*, and *spo-11 rec-8; coh-4 coh-3* mutant embryos. All bars = 4μm.

### Bipolar spindles assemble around single chromatids of metaphase I *spo-11 rec-8* embryos

To distinguish whether cohesin vs cohesion is required for bipolar spindle assembly, we analyzed *spo-11 rec-8* double mutants (Fig. 2C) which enter metaphase I with 24 single chromatids (Fig. 2D) but have been reported to retain COH- 3/4 cohesin on pachytene chromosomes (Severson et al., 2009). Bipolar metaphase I spindles assembled in *spo-11 rec-8* double mutants and these spindles shortened, rotated, and then segregated the chromatids into two masses (Fig. 2C, -6:50- 5:20; Video S4). At metaphase II, an amorphous mass of microtubules assembled around the chromatids and this mass shrank but did not separate chromatids into two masses (Fig. 2C, 16:10-18:40), similar to meiosis I in the quadruple mutant and meiosis II in both the quadruple mutant and the *rec-8* single mutant. The spindle pole protein, GFP::MEI-1, clearly labelled two poles of metaphase I and metaphase II control spindles but only labelled spindle poles of metaphase I *spo-11 rec-8* mutants (Fig. 2E). GFP::MEI-1 was dispersed on metaphase II spindles, confirming the apolar structure of these spindles. GFP::MEI-1 also associated with chromosomes and this chromosome association was much more apparent in metaphase II *spo-11 rec-8* spindles (Fig. 2E). However, the background subtracted ratio of mean GFP::MEI-1 pixel intensity on chromosomes divided by mean cytoplasmic intensity was not significantly increased between metaphase I and metaphase II for either *spo-11 rec-8* (MI: 7.01 ± 0.89, N=5 embryos, n=15 chromosomes; MII: 5.62 ± 0.76, N=5, n=15; p=0.23) or control spindles (MI: 5.62 ± 0.33, N=6, n=18; MII: 5.47 ± 0.35, N=6, n=18; p=0.74). This result indicated that the enhanced contrast of chromosomal GFP::MEI-1 in *rec-8 spo-11* embryos was due to the decrease in microtubule-associated GFP::MEI-1.

The ability of *spo-11 rec-8* embryos to form bipolar metaphase I spindles might be due to one or two univalents held together by residual COH-3/COH-4 cohesin.

However, 24 chromosome bodies could be counted in Z-stacks of the majority of metaphase I spindles (Fig. 2F) and all metaphase I spindles were bipolar (13/13 mNeonGreen tubulin, 9/9 GFP::MEI-1).

### Cohesin rather than cohesion is required for bipolar spindle assembly

The ability of *spo-11 rec-8* mutants to build bipolar metaphase I spindles but not metaphase II spindles might be because metaphase I chromatids retain cohesin, as high levels of COH-3/4 associate with pachytene chromosomes of *rec-8* mutants (Severson et al., 2009; Woglar et al., 2020). This non-cohesive COH-3/4 cohesin might be removed by separase at anaphase I, leaving the metaphase II chromatids with no cohesin. This hypothesis was validated by time-lapse imaging of the cohesin subunit, SMC- 1::AID::GFP, which would be a component of both REC-8 cohesin and COH-3/4 cohesin. SMC-1::AAID::GFP was found on control metaphase I and metaphase II chromosomes and metaphase I chromosomes of *spo-11 rec-8* mutants but was absent from the metaphase II chromatids of *spo-11 rec-8* mutants (Fig. 3A, 3B). To more directly test the requirement for cohesin, we monitored metaphase I spindle assembly in embryos depleted of SMC-1 with an auxin-induced degron (Castellano-Pozo et al., 2020). The majority of SMC-1-depleted embryos formed apolar metaphase I spindles (Fig. 3C). The small number of multipolar spindles likely resulted from an incomplete depletion of SMC-1. These results support the idea that cohesin on chromosomes rather than cohesion between chromosomes is required for bipolar spindle assembly during both meiosis I and meiosis II.

**Figure 3.**
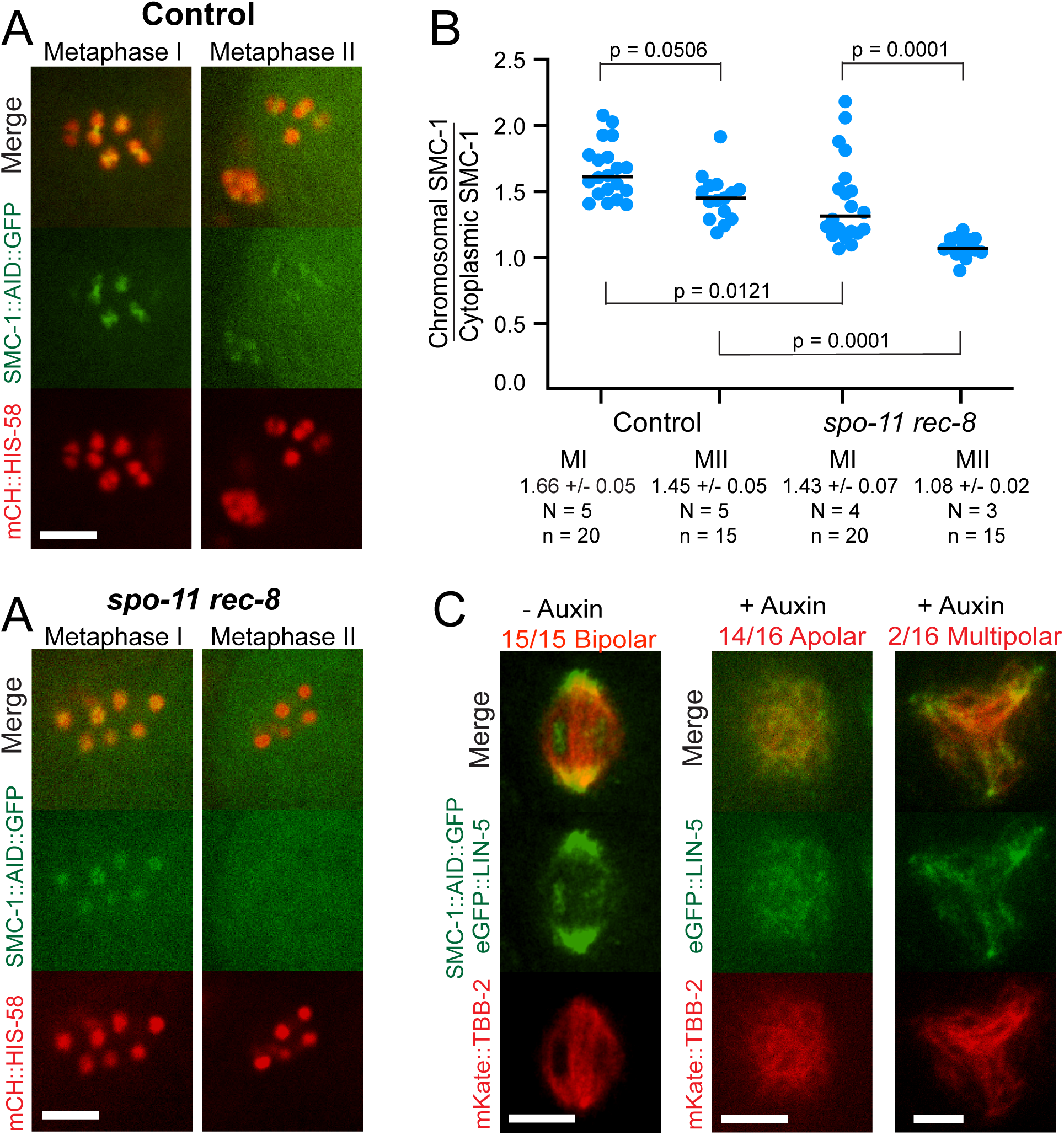
Non-cohesive cohesin is sufficient for bipolar spindle formation. **(A)** Single-plane images from control and *spo-11 rec-8* embryos expressing SMC-1::AID::GFP and mCH::HIS-58. **(B)** SMC-1::AID::GFP pixel intensities on individual chromosomes were determined relative to cytoplasmic background. N, number of embryos. n, number of chromosomes. **(C)** *C. elegans* expressing SMC-1::AID::GFP, eGFP::LIN-5 and mkate::TBB-2 were incubated overnight in the presence or absence of auxin. Single slices of z-stack MI images are shown. All bars = 4μm.

### A specific subclass of chromosome-associated Aurora B kinase correlates with competence for bipolar spindle assembly

We then asked why cohesin might be required for bipolar spindle assembly. In mitosis, cohesin-associated PDS5 recruits haspin kinase to chromosomes and the recruited haspin phosphorylates histone H3 threonine 3. The survivin (BIR-1 in *C. elegans*) subunit of the CPC binds to the phosphorylated histone thereby recruiting Aurora B to chromosomes (Kelly et al., 2010; Wang et al., 2010; Yamagishi, Honda, Tanno, & Watanabe, 2010). In *C. elegans*, haspin (HASP-1) is required to promote recruitment of Aurora B (AIR-2) to the midbivalent region in diakinesis oocytes (Ferrandiz et al., 2018) and AIR-2 is essential for bipolar meiotic spindle assembly in *C. elegans* (Divekar et al., 2021; Dumont et al., 2010), therefore we hypothesized that chromatids that lack cohesin-recruited AIR-2 would be unable to form bipolar meiotic spindles. Time-lapse imaging of control embryos with endogenously tagged AIR-2::GFP (Fig. 4A) revealed bright rings between homologs at metaphase I, microtubule association during anaphase I, bright rings between sister chromatids at metaphase II, and microtubule association during anaphase II as previously described (Dumont et al., 2010). In *rec-8* embryos, AIR-2 formed bright structures between sister chromatids at metaphase I and transferred to microtubules at anaphase I, as it does in controls. However, at metaphase II in *rec-8* embryos, AIR-2::GFP was dim and diffuse on spindle-incompetent single chromatids, then became bright on microtubules at anaphase II (Fig. 4B). In *rec-8* embryos, AIR- 2::GFP was significantly dimmer on chromosomes at metaphase II relative to metaphase I whereas no such decrease was observed in control embryos (Fig. 4C).

**Figure 4.**
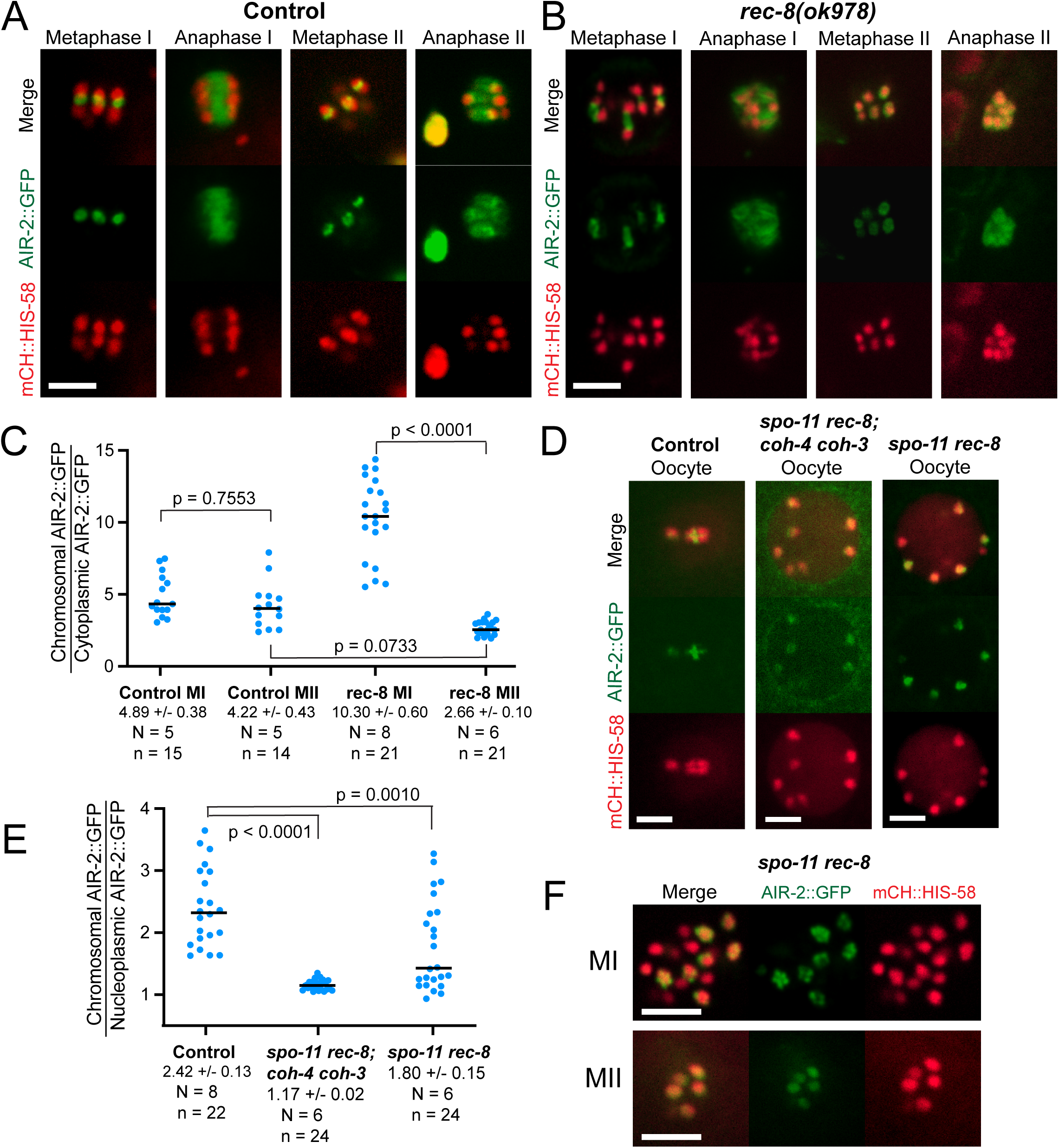
AIR-2::GFP levels are diminished and diffuse in the absence of cohesin. **(A)** In Control embryos, AIR-2::GFP is in the ring structure during metaphase I and II and on MTs during anaphase I and II. **(B)** In *rec-8(ok978)*, AIR-2::GFP is in the ring structure during MI, diffuse on chromatids in MII and on MTs during both anaphase I and II. **(C)** Quantification of AIR-2::GFP intensities on chromosomes relative to the cytoplasm in control and *rec-8(ok978)*. Ratios varied depending on the distance of the chromosomes from the objective. N, number of embryos. n, number of chromosomes. **(D)** -1 oocyte nuclei in control and mutant worms expresing AIR-2::GFP and mCH::HIS-58. **(E)** Quantification of AIR-2::GFP intensities on chromosomes relative to the nucleoplasm in control and mutant oocytes.. N, number of oocytes. n, number of chromosomes. **(F)** MI and MII metaphase chromosomes in *spo-11(me44) rec-8(ok978)* embryos. All bars = 4μm.

In control -1 diakinesis oocytes, which will initiate meiosis I spindle assembly within 1 – 23 min (McCarter, Bartlett, Dang, & Schedl, 1999), AIR-2::GFP brightly labeled the space between the homologous chromosomes in 6 bivalents. In contrast, GFP::AIR-2 was dim and diffuse on the spindle-incompetent single chromatids of *spo- 11 rec-8 coh-4 coh-3* quadruple mutants (Fig. 4D). Diakinesis oocytes of spindle- competent *spo-11 rec-8* double mutants contained a mixture of single chromatids with either dim diffuse AIR-2::GFP or bright patterned AIR-2::GFP (Fig. 4D, 4E). The bright patterned AIR-2::GFP on a subset of single chromatids could also be observed in bipolar metaphase I spindles of *spo-11 rec-8* mutants. In spindle-incompetent metaphase II embryos of *spo-11 rec-8* embryos, AIR-2::GFP was again dim and diffuse on all single chromatids (Fig. 4F). These results indicated that a specific subclass of AIR-2::GFP, that which is cohesin-dependent and forms a bright pattern on chromosomes, can promote bipolar spindle assembly. The subclasses of AIR-2::GFP that are cohesin-independent label chromatin dimly and diffusely, and label anaphase microtubules, but cannot promote bipolar spindle assembly. In support of this hypothesis, sperm-derived paternal DNA within meiotic embryos recruited maternal GFP::AIR-2 but lacked detectable cohesin and did not promote spindle assembly (Fig. S3). The cohesin-dependent subclass of AIR-2 might have a unique substrate specificity or it might be needed to reach a threshold of activity in combination with cohesin-independent AIR-2.

### Haspin-dependent Aurora B kinase is required for bipolar meiotic spindle assembly

To more specifically identify the subclass of Aurora B that is required for bipolar spindle assembly, we analyzed a *bir-1(E69A, D70A)* mutant. This double mutation is equivalent to the D70A, D71A mutation in human survivin that prevents binding to T3-phosphorylated histone H3 and prevents recruitment of Aurora B to mitotic centromeres in HeLa cells (Wang et al., 2010). Time-lapse imaging of mNeonGreen::tubulin in *bir-1(E69A, D70A)* mutants revealed apolar metaphase spindles that shrank without chromosome separation during both meiosis I and meiosis II (Fig. 5A). The *bir-1(E69A, D70A)* embryos were unlike the cohesin mutants in that they entered meiosis I with 6 bivalents (11/11 z-stacks of -1 oocytes), suggesting successful formation of chiasmata between homologous chromosomes during meiotic prophase and intact SCC (Figure 5A). Endogenously-tagged AIR-2::GFP diffusely labeled both lobes of metaphase I (Fig. 5B) and diakinesis (Fig. 5C) bivalents in *bir- 1(E69A, D70A)*. This was in contrast to the bright ring of AIR-2::GFP that is observed between the lobes in controls. AIR-2::GFP localized normally to microtubules during anaphase I and anaphase II (Fig. 5B) as was observed in cohesin mutants. Apolar metaphase I spindles (Fig. 5D, left) also formed after depletion of haspin kinase with an auxin-induced degron. Like *bir-1(E69A, D70A)* embryos, haspin-depleted embryos entered meiosis I with 6 bivalents (10/10 z-stacks of metaphase I), indicating the presence of chiasmata and SCC. As with cohesin mutants that were spindle- incompetent, the fluorescence intensity of AIR-2::GFP on chromosomes was strongly reduced in both *bir-1(E69A, D70A)* and *hasp-1(degron)* embryos (Fig. 5E). Whereas all *bir-1(E69A, D70A)* spindles were apolar, a minority of *hasp-1(degron)* spindles were multipolar (Fig 5D, Fig. 5F). This may be due to incomplete depletion by the degron.

**Figure 5.**
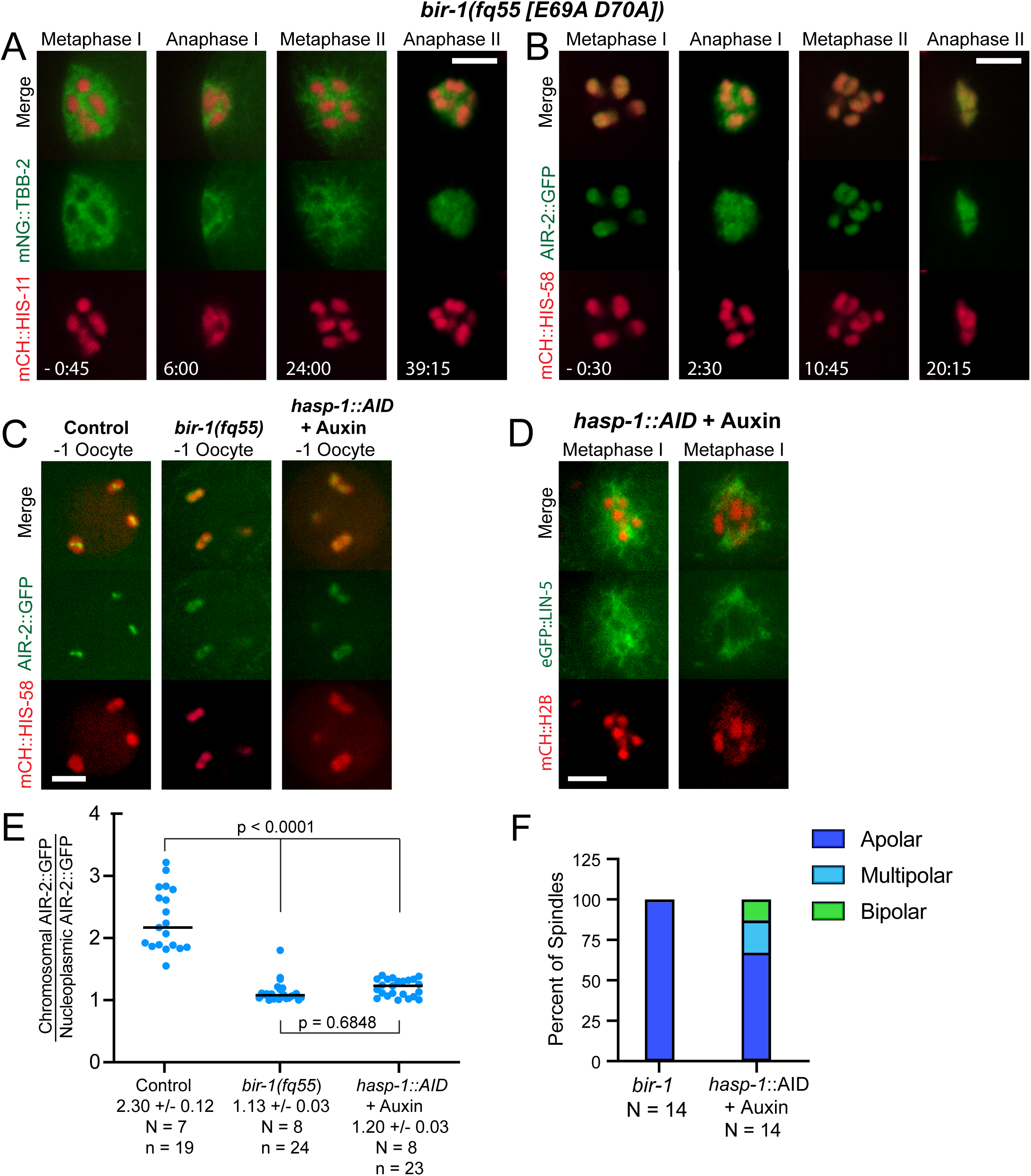
AIR-2 is recruited by Survivin and Haspin for bipolar spindle formation. **(A)** Time-lapse images of 5/5 *bir-1(fq55)* embryos expressing mNG::TBB-2 and mCH::HIS-11 show disorganized MI spindles and no MI anaphase chromosome separation. **(B)** Similar results were obtained in 4/4 *bir-1(fq55)* embryos expressing AIR-2::GFP, which is diffuse on both MI and MII metaphase chromosomes and present on MTs during anaphase. **(C)** Single slices from z-stack images of -1 oocytes in *C. elegans* expressing AIR-2::GFP and mCH::HIS-58. 11/11 -1 oocytes in *bir-1(fq55)* embryos had 6 mCH::HIS-58 labelled bodies. **(D)** Single-plane images of Auxin-treated *hasp-1::AID* embryos expressing eGFP::LIN-5 and mCH::H2B. 10/10 MI spindles in Auxin-treated *hasp-1:::AID* embryos had 6 mCH::HIS-58 labelled bodies. **(E)** AIR-2::GFP pixel intensities on individual chromosomes were determined relative to nucleoplasmic background. N, number of oocytes. n, number of chromosomes. **(F)** Graph showing percent of apolar, multipolar, and bipolar spindles in *bir-1* and auxin-treated *hasp-1::AID* embryos. N, number ofembryos. All bars = 4 μm

Because haspin is recruited to chromosomes by cohesin-associated PDS5 (Yamagishi et al., 2010), these results indicated that the subclass of Aurora B that is recruited to chromosomes by cohesin and haspin-dependent phosphorylation of histone H3 is required for bipolar spindle assembly and that cohesin-independent and haspin- independent Aurora B on chromosome lobes and anaphase microtubules are not sufficient to drive bipolar spindle assembly.

### Cohesin-dependent Aurora B kinase is required for coalescence of microtubule bundles into spindle poles

*C. elegans* meiotic spindle assembly begins at germinal vesicle breakdown in the -1 oocyte that is still part of the syncytial gonad. *In utero* time-lapse image sequences of spindle-competent control (Fig. 6A) and *spo-11 rec-8* (Fig. 6E) -1 oocytes, as well as spindle-incompetent *bir-1(E69A, D70A)* (Fig. 6C) and *spo-11 rec-8 coh-4 coh-3* (Fig. 6D) -1 oocytes, show that microtubule bundles initially filled the entire volume of the germinal vesicle as it broke down. Oocytes are then fertilized as they ovulate into the spermatheca. The microtubule bundles of control (Fig. 6B) and *spo-11 rec-8* (Fig. 6F) coalesced first into multiple poles, then into two poles as the oocytes squeezed into the spermatheca. In contrast, the microtubule bundles of *bir-1(E69A, D70A)* (Fig. 6C) and *spo-11 rec-8 coh-4 coh-3* (Fig. 6D) did not coalesce. This phenotype is consistent with that previously observed by fixed immunofluorescence of *air-2(degron)* embryos (Divekar et al., 2021). In addition, the mean fluorescence intensity of mNeonGreen::tubulin, indicative of microtubule density, was significantly reduced in apolar metaphase I spindles of *bir-1(E69A, D70A)* and *spo- 11 rec-8 coh-4 coh-3* embryos relative to the bipolar spindles in control and *spo-11 rec- 8* metaphase I spindles (Fig. S2). These results suggested that cohesin-dependent AIR-2 regulates proteins that coalesce microtubule bundles and promote microtubule polymerization.

### Cohesin-dependent Aurora B kinase regulates the localization of spindle assembly factors on meiotic chromosomes

We hypothesized that cohesin- dependent Aurora B on chromosomes might activate or inhibit microtubule-binding proteins that are required for coalescence of microtubule bundles or microtubule polymerization. Meiotic chromosome-associated spindle assembly factors include the katanin homolog, MEI-1 (Srayko, Buster, Bazirgan, McNally, & Mains, 2000), the kinesin-13, KLP-7 (Connolly, Sugioka, Chuang, Lowry, & Bowerman, 2015; Gigant et al., 2017), and the CLASP2 homolog, CLS-2 (Dumont et al., 2010; Schlientz & Bowerman, 2020). Loss of MEI-1 function results in apolar spindles with dispersed ASPM-1 (K. P. McNally & McNally, 2011) and reduced microtubule density (K. McNally, Audhya, Oegema, & McNally, 2006; Srayko, O’toole, Hyman, & Müller-Reichert, 2006) similar to those observed in cohesin mutants. However, apolar spindles in *mei-1* mutants are far from the cortex at metaphase I (Yang, McNally, & McNally, 2003) whereas cohesin-mutant apolar spindles were cortical at metaphase I (Fig. 1C, 1D, 2A, 2B). In addition, endogenously tagged GFP::MEI-1 was retained on chromosomes of apolar metaphase II *spo-11 rec-8* mutants (Fig. 2E). These results suggest that MEI-1 is active in embryos that are deficient in cohesin-recruited AIR-2.

**Figure 6.**
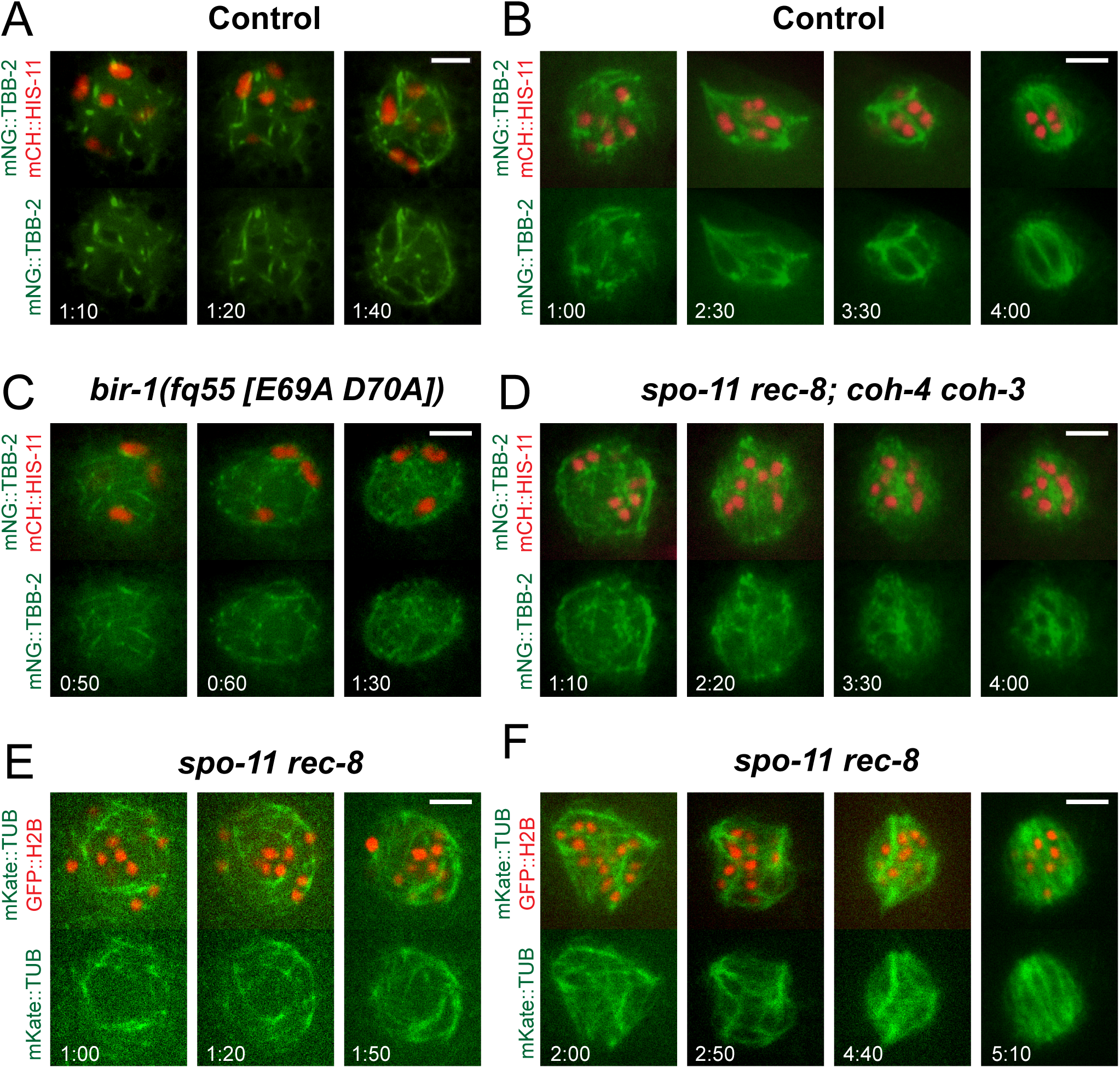
Cohesin-recruited AIR-2 is necessary to direct the formation of organized spindle fibers. (A, B) Time-lapse images of 7/7 Control embryos show MT fibers organizing rapidly around chromosomes. Spindles become multipolar, then bipolar as poles coalesce. Times are from the initial observation of spindle MTs. **(C)** Time-lapse images of 7/7 *bir-1* embryos and **(D)** 7/7 *spo-11 rec-8; coh-4 coh-3* embryos show that spindle fibers begin to form, but do not become organized. **(E, F)** Time-lapse images of 13/13 *spo-11 rec-8* embryos show spindle fibers rapidly engulfing chromosomes. Images in E and F have been pseudocolored for increased clarity. All bars = 4μm.

Endogenously tagged KLP-7::mNeonGreen localized to the midbivalent ring and to the two lobes of control bivalents but localized only to the two lobes in *bir-1(E69A, D70A)* mutants (Fig. 7A). In *spo-11 rec-8* double mutants, KLP-7::mNeonGreen localized in a bright pattern with a larger area on a subset of single chromatids in spindle-competent metaphase I embryos but labeled single chromatids with a more uniform smaller area in spindle-incompetent metaphase II embryos (Fig. 6B, C). In *spo- 11 rec-8* metaphase I embryos there was a positive correlation between the fluorescence intensity of endogenously tagged mScarlet::AIR-2 and the area of endogenously tagged KLP-7::mNeonGreen (Fig. 6D, E). This result indicated that the subclass of AIR-2 that can promote bipolar spindle assembly also recruits a subclass of KLP-7 to chromosomes.

**Figure 7.**
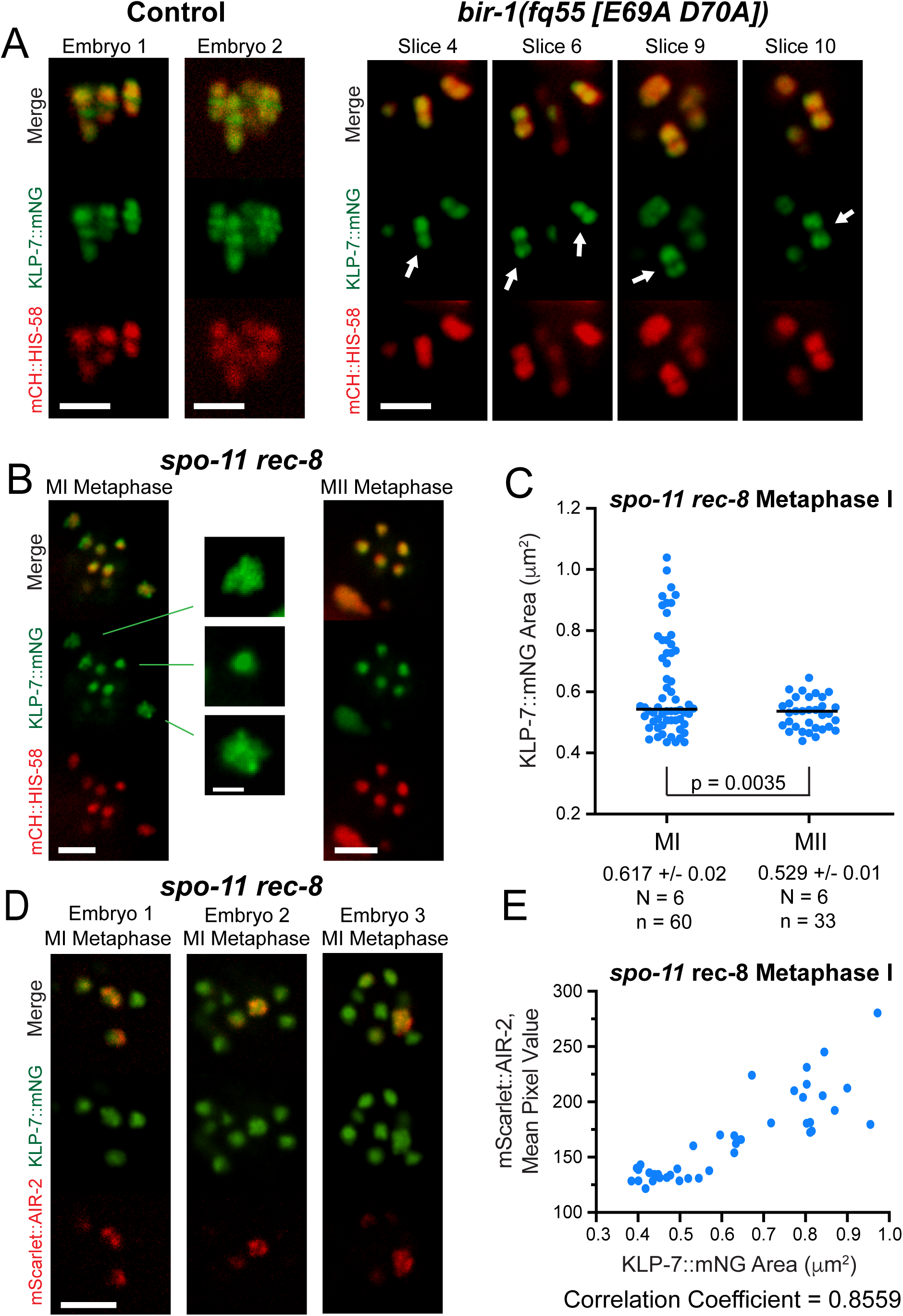
Survivin-dependent AIR-2 is required for KLP-7 recruitment to the ring complex. (A and B) KLP-7::mNG localizes to the centromere and ring complex in 7/7 control embryos, but is only centromeric in 8/8 *bir-1(fq55) [E69A D70A]) embryos.* **(B)** In *spo-11 rec-8* MI metaphase spindles, one subset of chromosomes has only centromeric KLP-7::mNG and a second subset has KLP-7::mNG dispersed over an expanded area. Bar, spindle images, 3 μm. Bar, single chromosome images, 1 μm. **(C)** KLP-7::mNG areas were determined in *spo-11 rec-8* MI metaphase and MII metaphase spindles. N, number of embryos. n, number of chromosomes. **(D)** Single z-stack images of 14/14 *spo-11* rec-8 embryos show expanded KLP-7::mNG on chromosomes with highest wrmScarlet::AIR-2 levels. Bar, 3 μm. **(E)** Graph of wrmScarlet::AIR-2 mean pixel value relative to KLP-7::mNG area. The Pearson r correlation coefficient is 0.8559.

CLS-2::GFP labeled the kinetochore cups enveloping the two lobes of metaphase I bivalents but was excluded from the midbivalent ring in control embryos. In contrast, CLS-2::GFP labeled kinetochore cups and the midbivalent ring in *bir- 1(E69A, D70A)* mutants (Fig. 8A). In *spo-11 rec-8* double mutants, CLS-2::GFP localized in spheres with a larger diameter on a subset of single chromatids in spindle- competent metaphase I embryos but labeled single chromatids with a more uniform smaller diameter in spindle-incompetent metaphase II embryos (Fig. 7B, C). In *spo-11 rec-8* metaphase I embryos there was a positive correlation between the fluorescence intensity of endogenously tagged mScarlet::AIR-2 and the diameter of CLS-2::GFP spheres (Fig. 7D, E). These results indicate that cohesin-dependent AIR-2 both excludes CLS-2 from the midbivalent ring and recruits CLS-2 into larger spheres around single chromatids.

**Figure 8.**
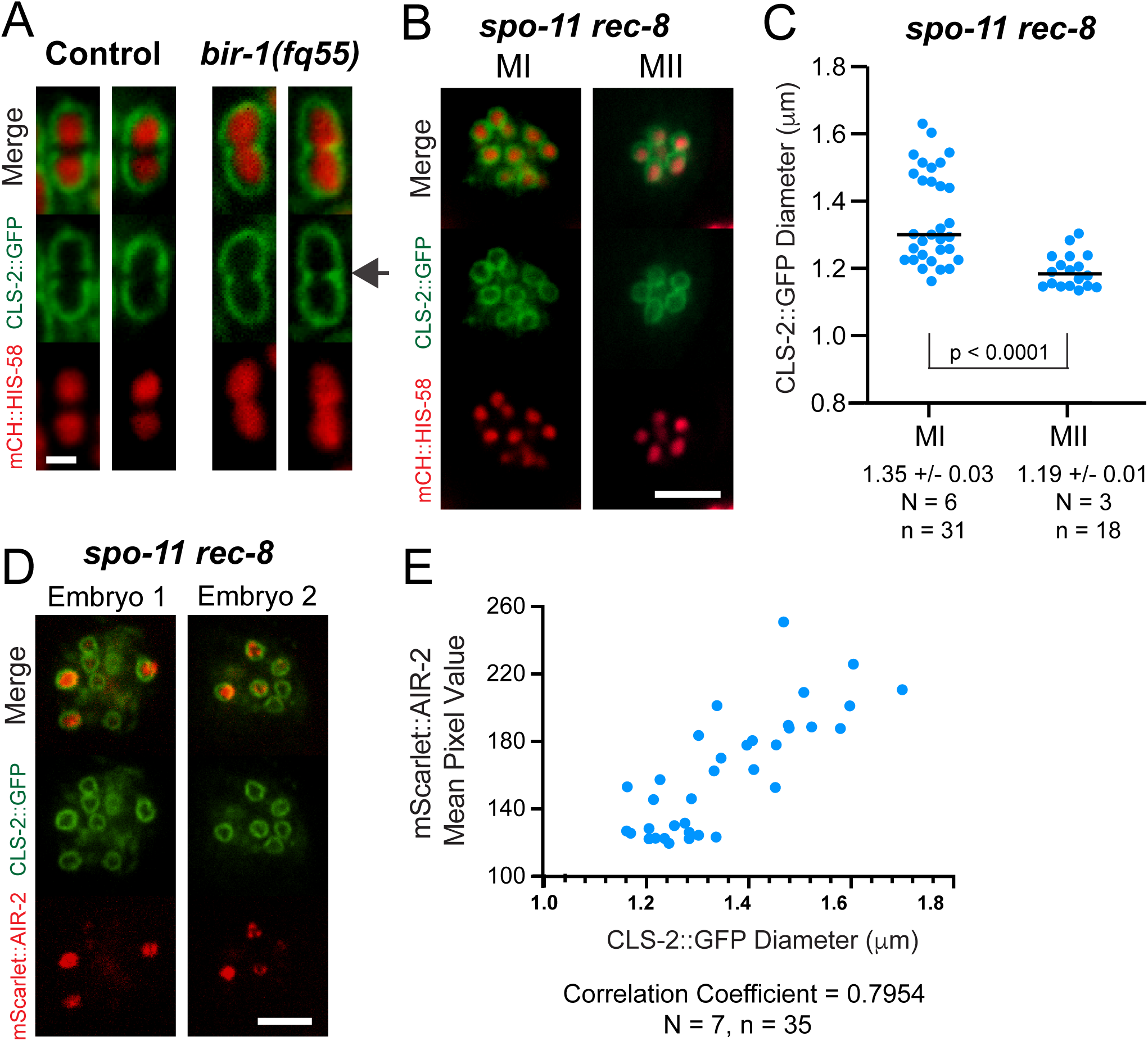
BIR-1-recruited AIR-2 excludes CLS-2 from the ring structure. **(A)** Individual chromosomes in embryos expressing CLS-2::GFP show that CLS-2 is excluded from the ring structure in 9/9 control embryos and present in the ring structure in 16/16 *bir-1(fq55)* embryos. Bar = 1μm. **(B)** CLS-2::GFP encirles both MI and MII metaphase chromosomes in *spo-11 rec-8* embryos. **(C)** The diameter of CLS-2::GFP spheres on MI and MII metaphase chromosomes was determined. N, number of embryos. n, number of chromosomes. **(D)** Single z-stack slices of two *spo-11 rec-8* embryos expressing CLS-2::GFP and wrmScarlet::AIR-2. (**E)** Graph showing mean pixel value of wrmScarlet::AIR-2 versus CLS-2::GFP diameter on *rec-8; spo-11* chromosomes. Bars, B and D, 4μm. N, number of embryos. n, number of chromosomes.

### Most *C. elegans* meiotic SAFs are cytoplasmic

RNAi depletions of ran-GTP pathway components had relatively mild effects on meiotic spindle assembly in *C. elegans* (Chuang et al., 2020) however this could be due to incomplete depletion. Vertebrate ran-dependent SAFs bind importins and are nuclear during interphase (Cavazza & Vernos, 2015). To address the relative roles of the ran pathway and the CPC pathway, we imaged endogenously GFP-tagged SAFs in diakinesis oocytes just before initiation of spindle assembly (-1 oocytes). MEI-1, LIN-5, CLS-2, and AIR-1, which all contribute to bipolar spindle assembly in *C. elegans* (Dumont et al., 2010; Mains, Kemphues, Sprunger, Sulston, & Wood, 1990; Sumiyoshi, Fukata, Namai, & Sugimoto, 2015; van der Voet et al., 2009), were all cytoplasmic before nuclear envelope breakdown (Fig. S3). In addition, KLP-15/16, which are required for spindle assembly, have been reported to be cytoplasmic in -1 oocytes (Mullen & Wignall, 2017). These results indicate that most of the known SAFs during *C. elegans* meiosis cannot be activated by the canonical ran pathway.

## Discussion

Our results indicate that cohesin is required for acentrosomal spindle assembly independent of its role in SCC because it is required for recruitment of a specific pool of Aurora B kinase to chromatin. The requirement for cohesin is independent of SCC because single chromatids bearing COH-3/4 cohesin in *spo-11 rec-8* double mutants support the assembly of bipolar spindles. In contrast, single chromatids in mutants lacking any cohesin assembled amorphous masses of microtubules with no discrete foci of spindle pole proteins. The cohesin-dependent pool of Aurora B kinase is then required for microtubule bundles to coalesce to form spindle poles during *C. elegans* oocyte meiotic spindle assembly. In the absence of either cohesin, haspin kinase, or phosphorylated histone H3-bound survivin, Aurora B remains dispersed on metaphase chromatin and localizes on anaphase microtubules but is insufficient to promote spindle pole formation. This could be due to a need for a threshold concentration of Aurora B on chromatin or a need for a specific activity unique to cohesin-dependent Aurora B. Because separase normally removes residual cohesin at anaphase II, the cohesin-dependence of acentrosomal spindle assembly may help to ensure that a metaphase III spindle does not form before the first mitotic cell cycle .

The mechanism by which a specific pool of Aurora B kinase promotes spindle pole formation is not clear. Our studies suggest that Aurora B is required during a very early stage of spindle formation: the coalescence of MT bundles. Once formed, the MT bundles in *spo-11 rec-8* mutants appeared to self-assemble into a bipolar structure.

Completely apolar meiotic spindles have been observed in *C. elegans* upon depletion of MEI-1/2 katanin (K. P. McNally & McNally, 2011), KLP-15/16 (Mullen & Wignall, 2017), and AIR-1 (Sumiyoshi et al., 2015). If Aurora B inhibits a target SAF through phosphorylation, then the net effect of CPC loss would likely be hyperactivation rather than depletion of a SAF. Over-expression phenotypes have not been reported for *C. elegans* meiotic SAFs due to technical limitations of transgene technology. Loss of haspin-dependent CPC in this study caused a change in the localization pattern of KLP- 7 and CLS-2 on chromosomes. Loss of KLP-7 (Connolly et al., 2015; Gigant et al., 2017) or CLS-2 (Schlientz & Bowerman, 2020) results in multi-polar rather than apolar spindles but the effects of their over-expression or incorrect localization are not known. In *Xenopus* extracts, the primary function of Aurora B in spindle assembly has been suggested to be inhibition of the KLP-7 homolog, MCAK (Sampath et al., 2004).

Katanin is also inhibited in *Xenopus laevis* egg extracts by phosphorylation of an Aurora consensus site (Loughlin, Wilbur, McNally, Nedelec, & Heald, 2011). Whereas this exact site is not conserved in *C. elegans* MEI-1, the activity of MEI-1 is inhibited by phosphorylation at several sites (Joly, Beaumale, Van Hove, Martino, & Pintard, 2020). These results suggest that the overall effect of the CPC on spindle pole assembly could involve activation or inhibition of multiple SAFs that would be difficult to replicate with phosphorylation site mutants.

Similar to *C. elegans spo-11 rec-8* mutants, *Drosophila sunn* mutants lack SCC but retain non-cohesive c(2)m/SA cohesin (at least in pachytene) and form bipolar metaphase I spindles (Gyuricza et al., 2016). Depletion of SMC3, which should remove all cohesin from chromatin, has been reported in mouse oocytes (Yueh, Singh, & Gerton, 2021) and *Drosophila* oocytes (Gyuricza et al., 2016). Metaphase I spindle defects were not reported in either case. In both cases, cohesin depletion may have been incomplete, cohesin-independent Aurora B might suffice for spindle assembly, or the GTP-ran pathway might dominate in these species. Thus it remains unclear whether the cohesin-dependence of acentrosomal spindle assembly applies in phyla other than Nematoda.

Our time-lapse imaging revealed single chromatids separating into two masses during anaphase I in *spo-11 rec-8* embryos. This result is consistent with the previously published observation of a single polar body and equational segregation interpreted from polymorphism analysis (Severson et al., 2009). Similarly, *Drosophila sunn* mutants are able to carry out anaphase I (Gyuricza et al., 2016). HeLa cells induced to enter mitosis with unreplicated genomes likely have G1 non-cohesive cohesin on their single chromatids. These cells assemble bipolar spindles but do not separate the single chromatids into two masses. Instead, all of the chromatids end up in one daughter cell at cytokinesis (O’Connell et al., 2008). In *C. elegans* meiosis, anaphase B occurs by CLS-2-dependent microtubule pushing on the inner faces of separating chromosomes (Laband et al., 2017). During normal meiosis, the CPC recruits CLS-2 to the midbivalent ring (Dumont et al., 2010) so that microtubules push the correct homologous chromosomes apart. In a *spo-11 rec-8* double mutant, bright patterned AIR-2 is only on a subset of chromatids but microtubules still appeared to push all of the chromatids apart. Presumably, microtubules are pushing between any two chromatids. This *faux* anaphase likely occurs by the same mechanism as anaphase B in embryos depleted of outer kinetochore proteins (Danlasky et al., 2020; Dumont et al., 2010).

The spindle-competent single chromatids of *C. elegans spo-11 rec-8* mutants had a severe congression defect (Fig. 2C, D). In contrast, unreplicated single chromatids in HeLa cells congress normally to the metaphase plate (O’Connell et al., 2008). It is likely that antagonism between dynein in kinetochore cups and KLP-19 in the midbivalent ring is important for chromosome congression in *C. elegans* oocytes (Muscat, Torre-Santiago, Tran, Powers, & Wignall, 2015), thus the striking bipolar structure of *C. elegans* metaphase I bivalents and metaphase II univalents is essential for congression while dispensable for bipolar spindle assembly or anaphase.

## Materials and Methods

CRISPR-mediated genome editing to create the *bir-1(fq55[E69A D70A])* allele was performed by microinjecting preassembled Cas9sgRNA complexes, single-stranded DNA oligos as repair templates, and dpy-10 as a co-injection marker into the *C. elegans* germline as described in Paix et al (Paix, Folkmann, & Seydoux, 2017). The TCGTACCACGGATCGTCTTC sequence was used for the guide RNA and the single-stranded DNA oligo repair template had the following sequence: tgtgcattttgcaacaaggaacttgattttgaccccgctgctgacccgtggtacgagcacacgaaacgtgatgaaccgtg. *C. elegans* strains were generated by standard genetic crosses, and genotypes were confirmed by PCR. Genotypes of all strains are listed in Table S1.

### Live *in utero* Imaging

L4 larvae were incubated at 20°C overnight on MYOB plates seeded with OP50. Worms were anesthetized by picking adult hermaphrodites into a solution of 0.1% tricaine, 0.01% tetramisole in PBS in a watch glass for 30 min as described in Kirby et al. (Kirby, Kusch, & Kemphues, 1990) and McCarter et al. (McCarter et al., 1999).

Worms were then transferred in a small volume to a thin agarose pad (2% in water) on a slide. Additional PBS was pipetted around the edges of the agarose pad, and a 22-×- 30-mm cover glass was placed on top. The slide was inverted and placed on the stage of an inverted microscope. Meiotic embryos or -1 diakinesis oocytes were identified by bright-field microscopy before initiating time-lapse fluorescence. For all live imaging, the stage and immersion oil temperature was 22°C–24°C. For all live imaging data, single– focal plane time-lapse images were acquired with a Solamere spinning disk confocal microscope equipped with an Olympus IX-70 stand, Yokogawa CSU10, Hamamatsu ORCA FLASH 4.0 CMOS (complementary metal oxide semiconductor) detector, Olympus 100×/1.35 objective, 100-mW Coherent Obis lasers set at 30% power, and MicroManager software control. Pixel size was 65 nm. Exposures were 300 ms. Time interval between image pairs was 15 s with the exception of Figure 6 images, which were captured at 10 s intervals. Focus was adjusted manually during time-lapse imaging.

### Timing

Control spindles maintain a steady-state length of 8 µm for 7 min before initiating APC-dependent spindle shortening, followed by spindle rotation and movement to the cortex (Yang, Mains, & McNally, 2005). Because the majority of our videos began after MI metaphase onset, we measured time relative to the arrival of the spindle at the cortex in Figures 1, 2, and 5. For Figure 6, time was measured relative to the initial appearance of MT fibers.

### Auxin

*C. elegans* strains endogenously tagged with auxin-inducible degrons and a TIR1 transgene were treated with 4 mm auxin overnight on seeded plates.

### Fluorescence Intensity Measurements

Fluorescence intensity measurements are from single focal plane images. In Figures 3B, 4C 4E, and 5E, total pixel values of chromosomal SMC-1::AID::GFP or AIR- 2::GFP were obtained using the Freehand Tool (ImageJ software) to outline individual chromosomes. For each chromosome, the ROI was dragged to the adjacent nucleoplasm or cytoplasm and the total pixel value obtained. The values were background-subtracted and divided in order to generate a ratio for comparison. In Figure 7C and 7E, areas of KLP-7::mNG on individual chromosomes was measured using the Freehand Tool (ImageJ). The diameter of CLS-2::GFP rings in Figure 8 was calculated from the area, which was obtained by drawing an ellipse over each ring.

Mean mScarlet::AIR-2 pixel values in Figures 7 and 8 were determined after outlining individual chromosomes with the Freehand Tool (ImageJ). In Supplemental Figure 1, single-plane images were captured at the midsection of -1 oocytes. For each image, regions of nucleoplasm and cytoplasm were outlined and the mean pixel values determined. In Supplemental Figure 2, single-plane images were captured at the midsection of metaphase I spindles. For each image, mean pixel values of the spindle and a region of cytoplasm were determined. For both figures, the mean values were background-subtracted and divided to generate ratios for comparison.

### Statistics

P values were calculated in GraphPad Prism using one-way ANOVA for comparing means of three or more groups. Pearson correlation coefficients were calculated using GraphPad Prism.

## Acknowledgements

We thank Fede Pelisch, Arshad Desai, and the CGC, which is funded by NIH Office of Research Infrastructure Programs (P40 OD010440), for strains. This work was funded by grant R35GM136241 from the National Institute of general Medical Sciences and Hatch grant 1009162 from the U.S. Department of Agriculture/National Institute of Food and Agriculture to F.J.M and by MRC core-funded grant MC-A652-5PY60 to E.M.-P.

---

**Table S1.**
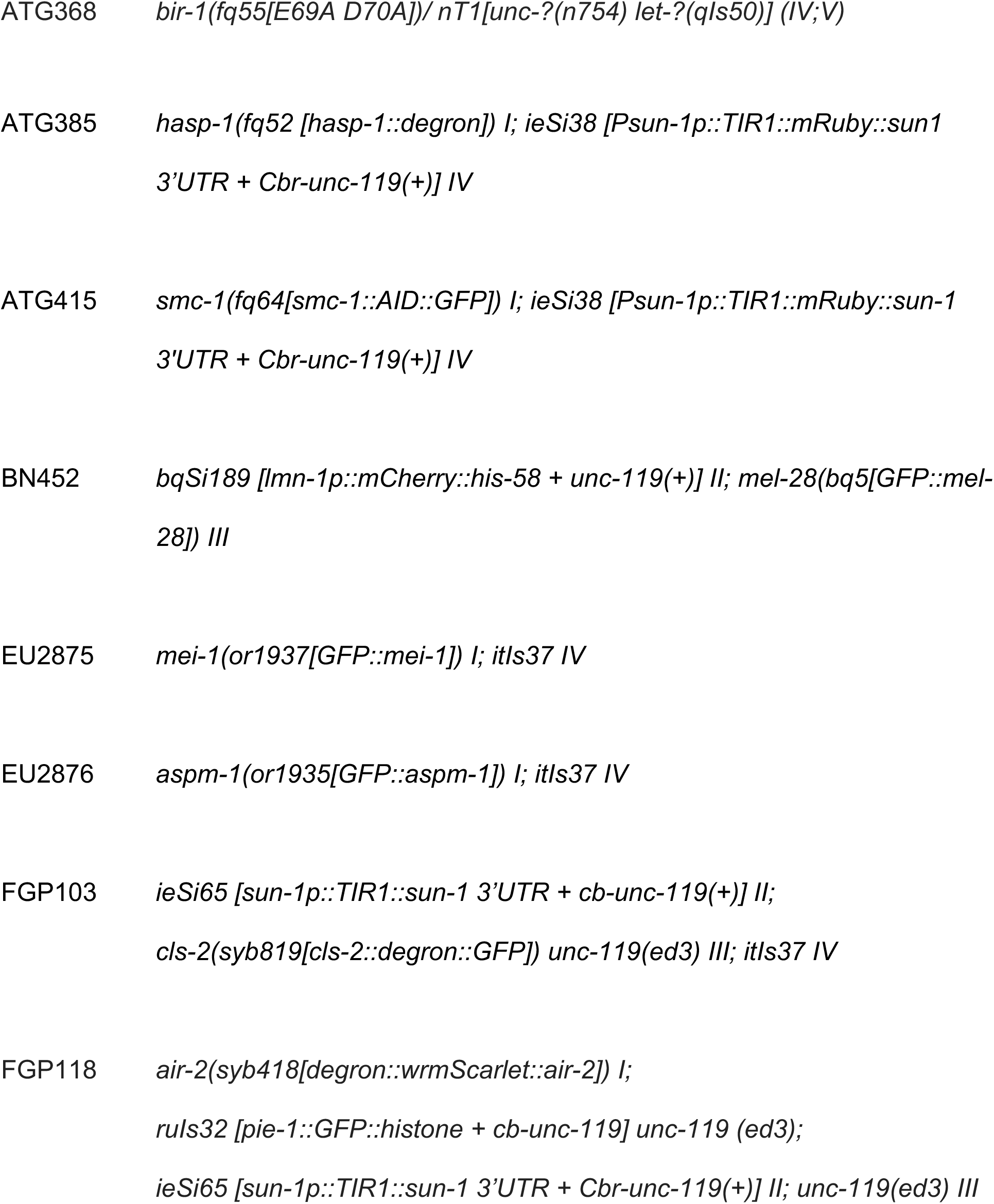
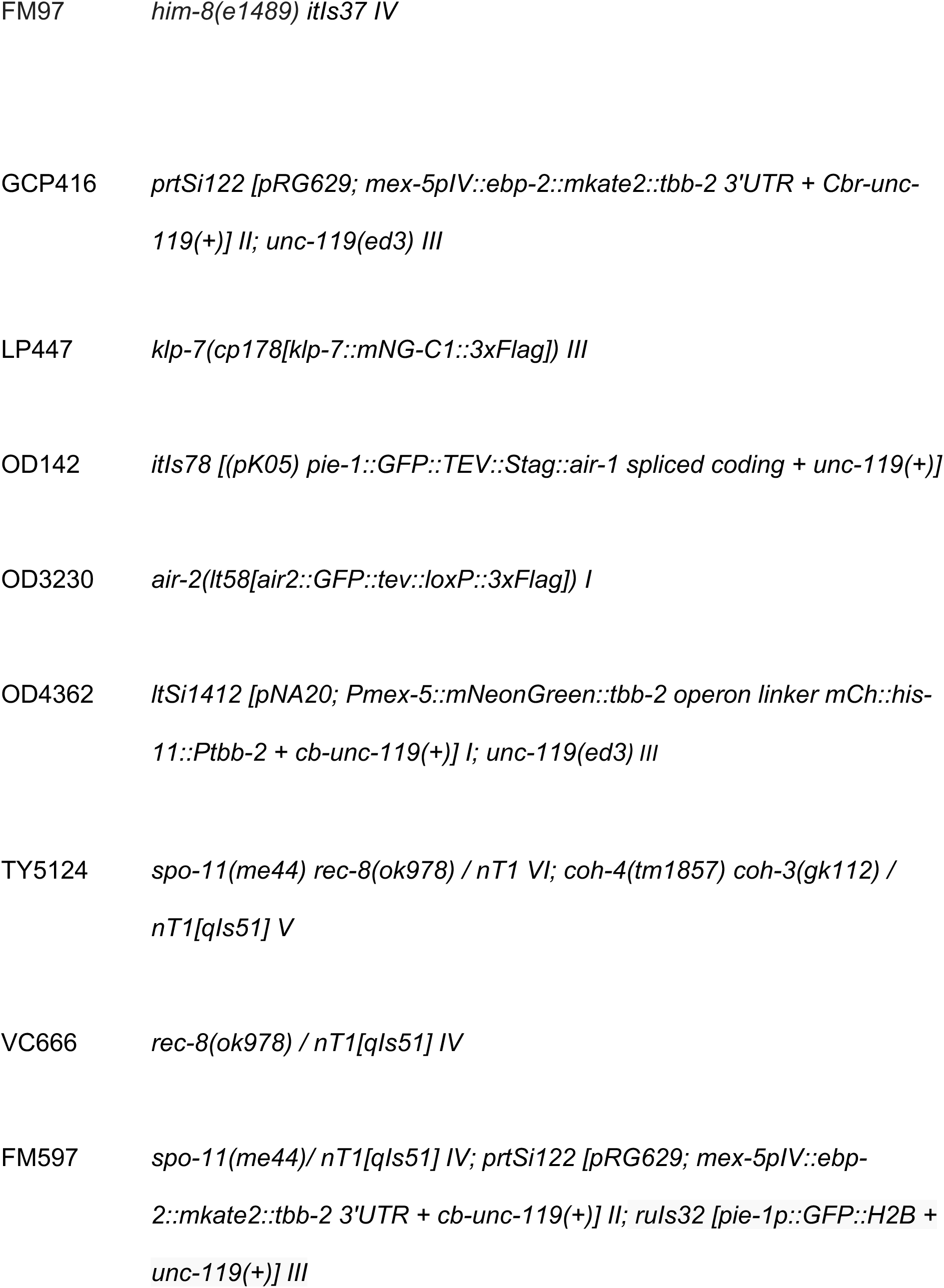
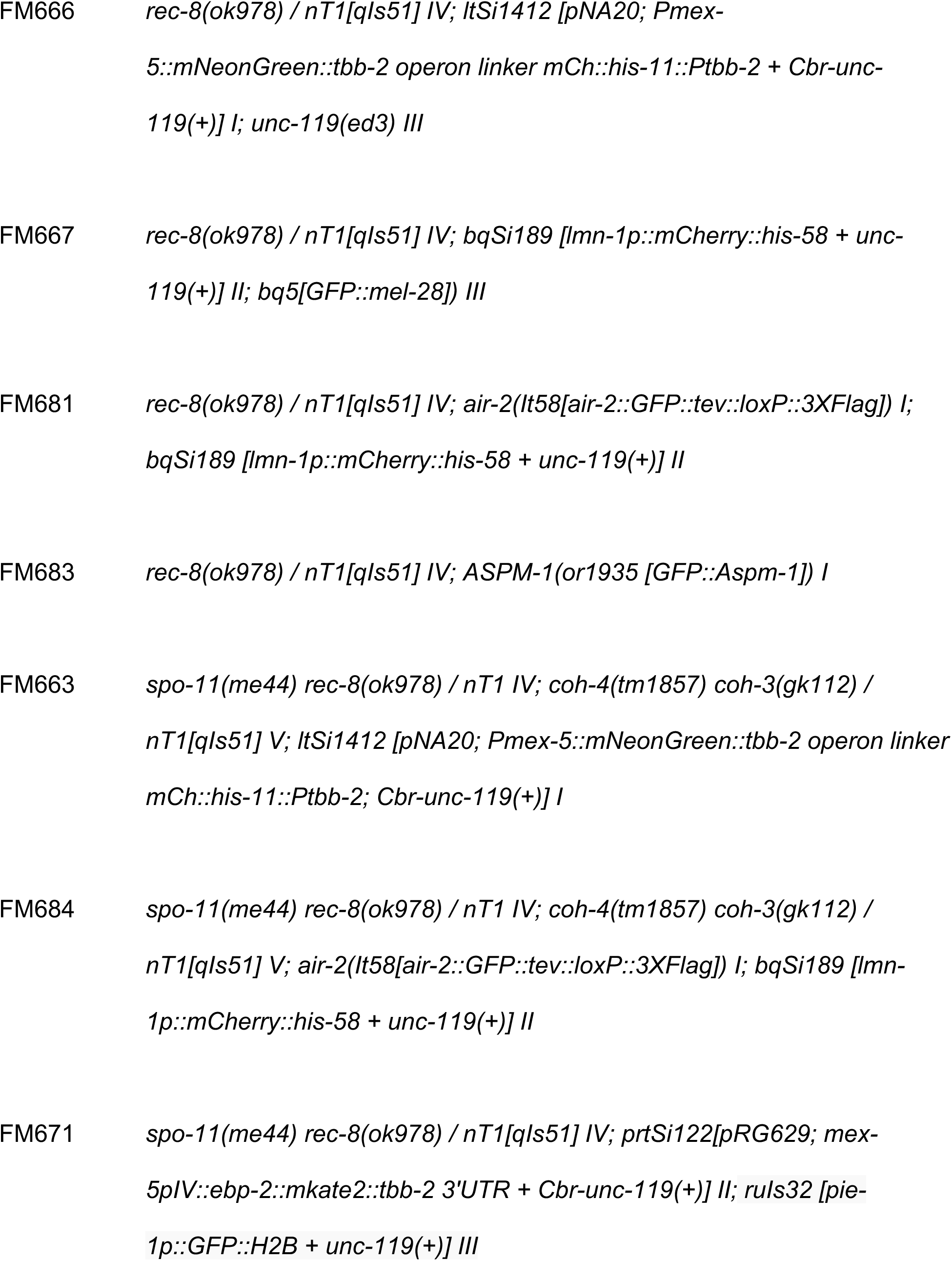
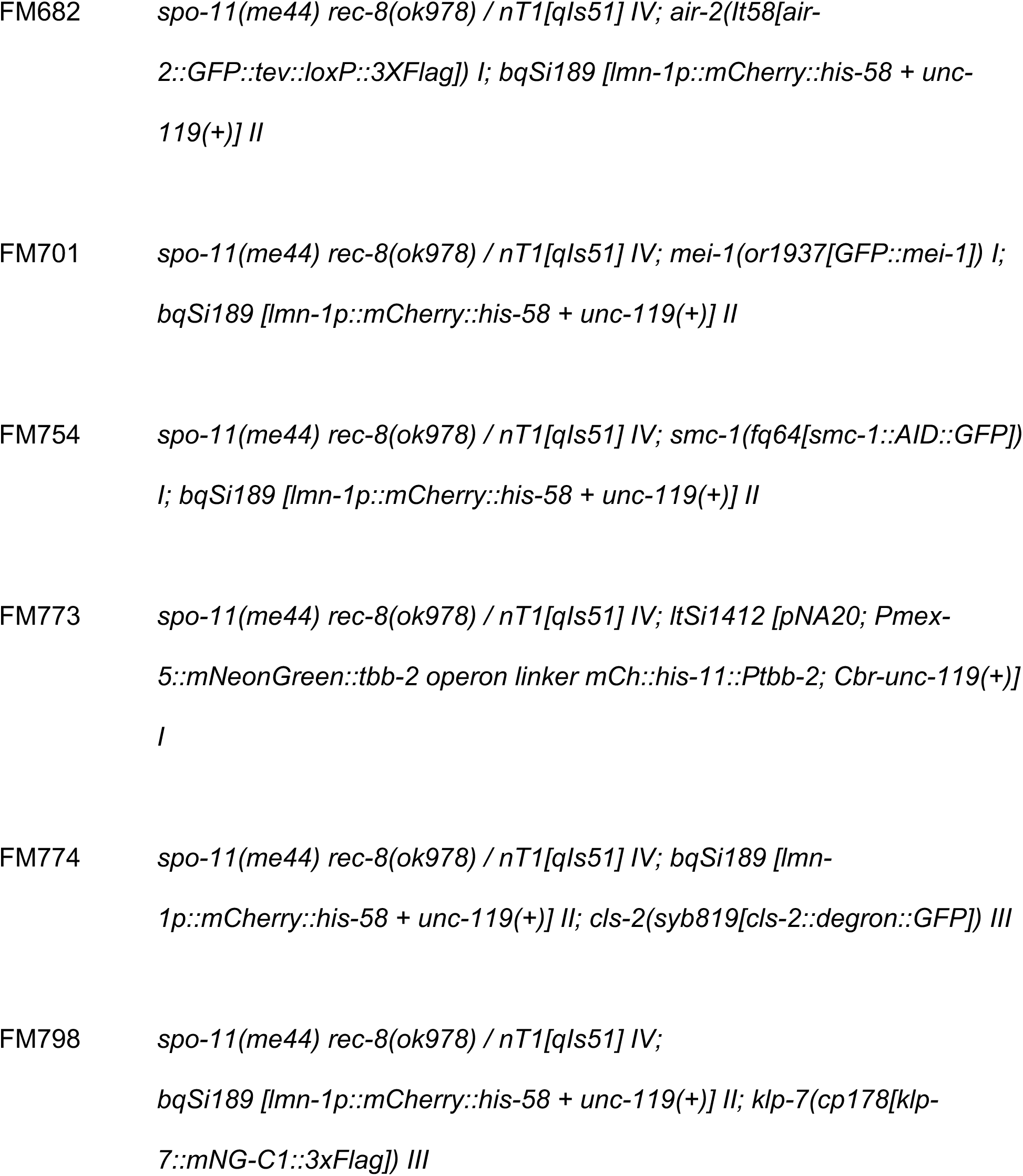
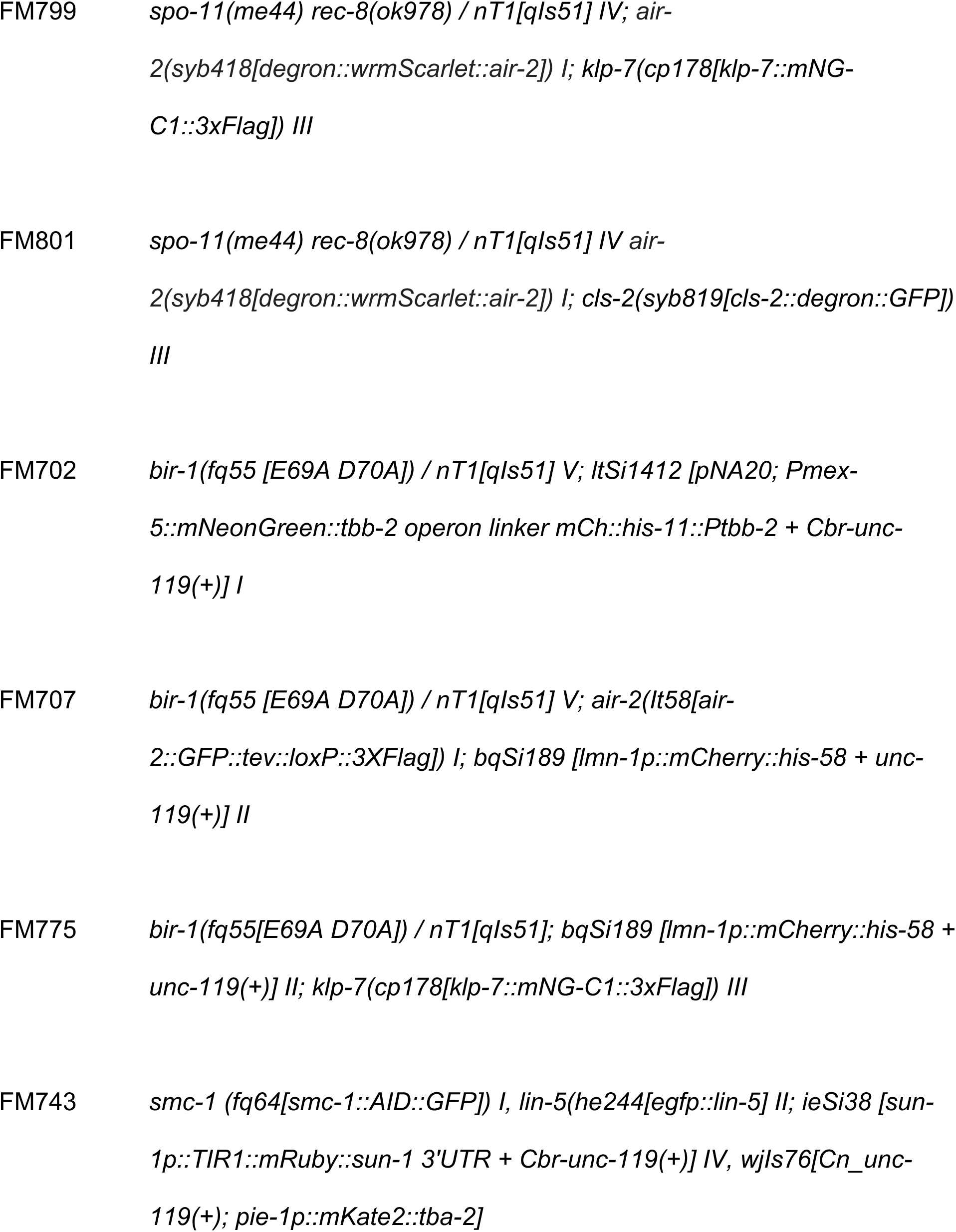
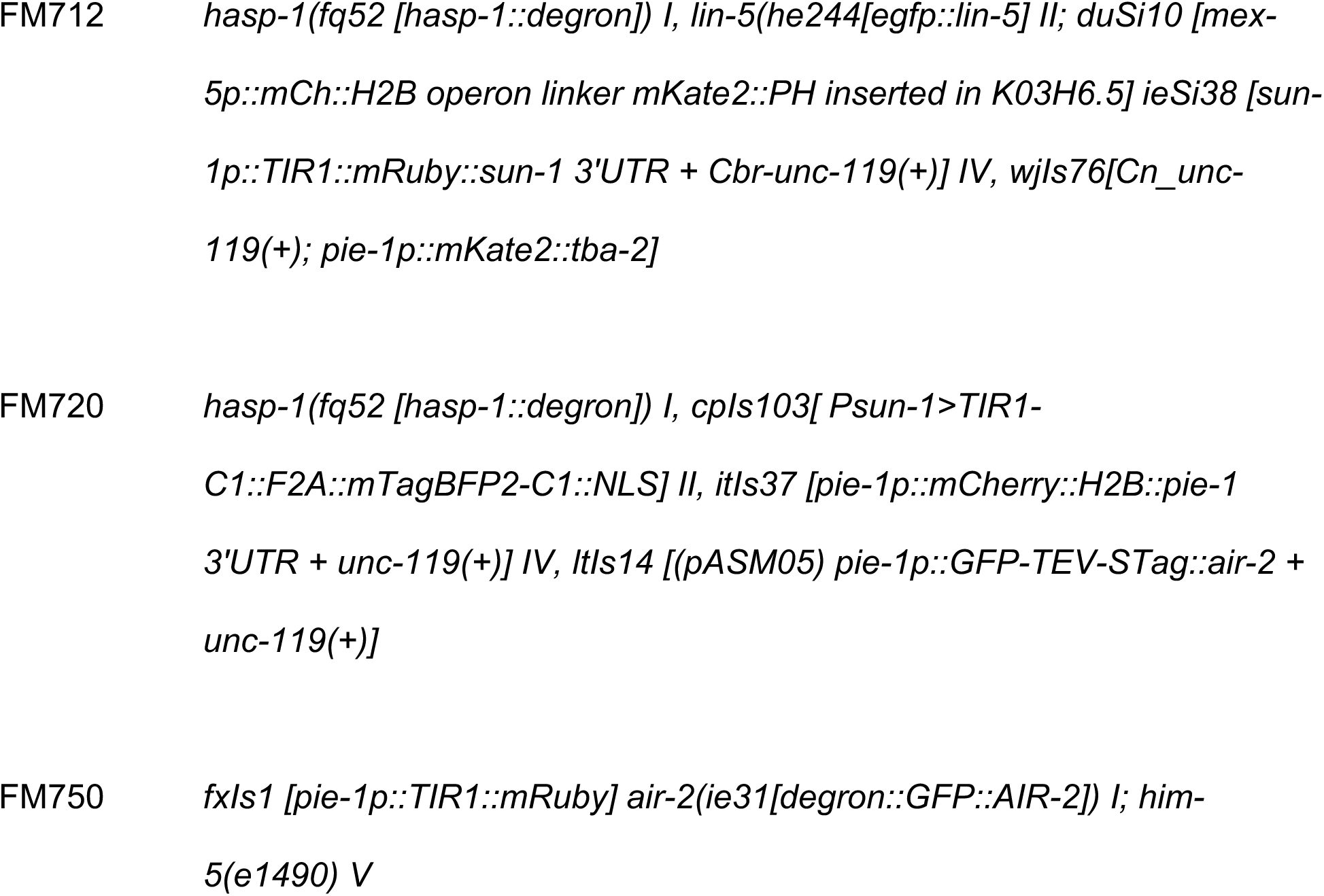
*C. elegans* Strain List

## Supplemental Video Legends

**Video S1**: Metaphase I through anaphase II filmed in utero in a control strain. Green is mNeonGreen::tubulin. Red is mCherry::histone H2b.

**Video S2**: Metaphase I through anaphase II filmed in utero in a *rec-8(ok978)* strain. Green is mNeonGreen::tubulin. Red is mCherry::histone H2b.

**Video S3**: Metaphase I through anaphase II filmed in utero in a *rec-8(ok978)* strain. Green is mNeonGreen::tubulin. Red is mCherry::histone H2b.

**Video S4**: Metaphase I through anaphase II filmed in utero in a *spo-11(me44) rec- 8(ok978)* strain. Green is GFP::histone H2b. Red is mKate::tubulin.

---

**Supplemental Figure 1.**
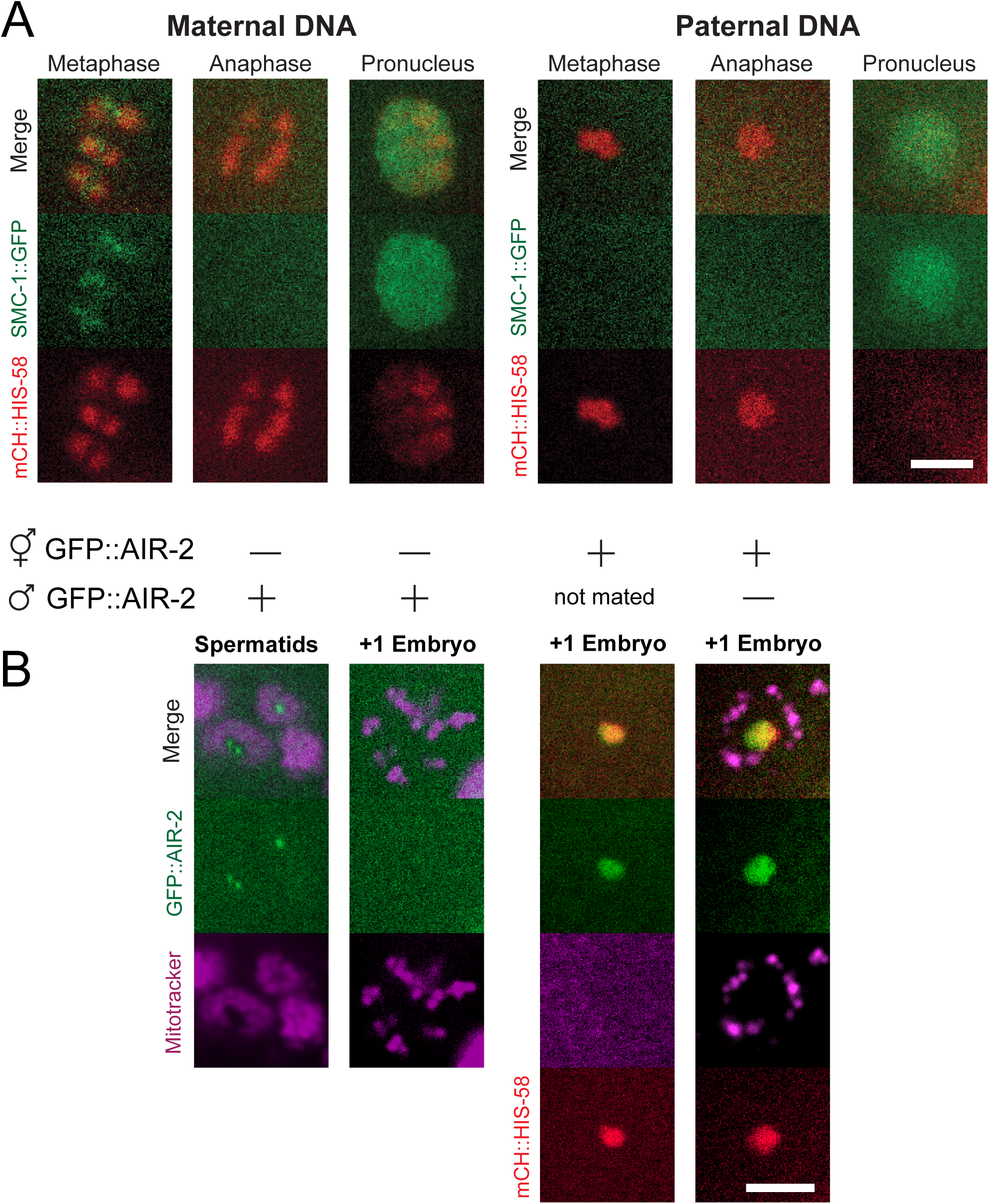
Maternal AIR-2, but not SMC-1, is recruited to the sperm DNA. **(A)** Time-lapse images of 15/15 embryos expressing SMC-1::GFP and mCH::HIS-58 show no SMC-1::GFP on sperm DNA during meiosis. SMC-1::GFP was observed in the paternal pronucleus in 7/7 embryos. Bar = 3 μm. (B) In 5/5 mated hermaphrodites, paternal GFP::AIR-2 is present on spermatids, but is not present post-fertilization. 13/13 unmated hermaphrodites expressing GFP::AIR-2, and 11/11 expressing hermaphrodites mated with non-expressing males have GFP::AIR-2 on the sperm DNA in +1 embryos. Bar = 4μm.

**Supplemental Figure 2.**
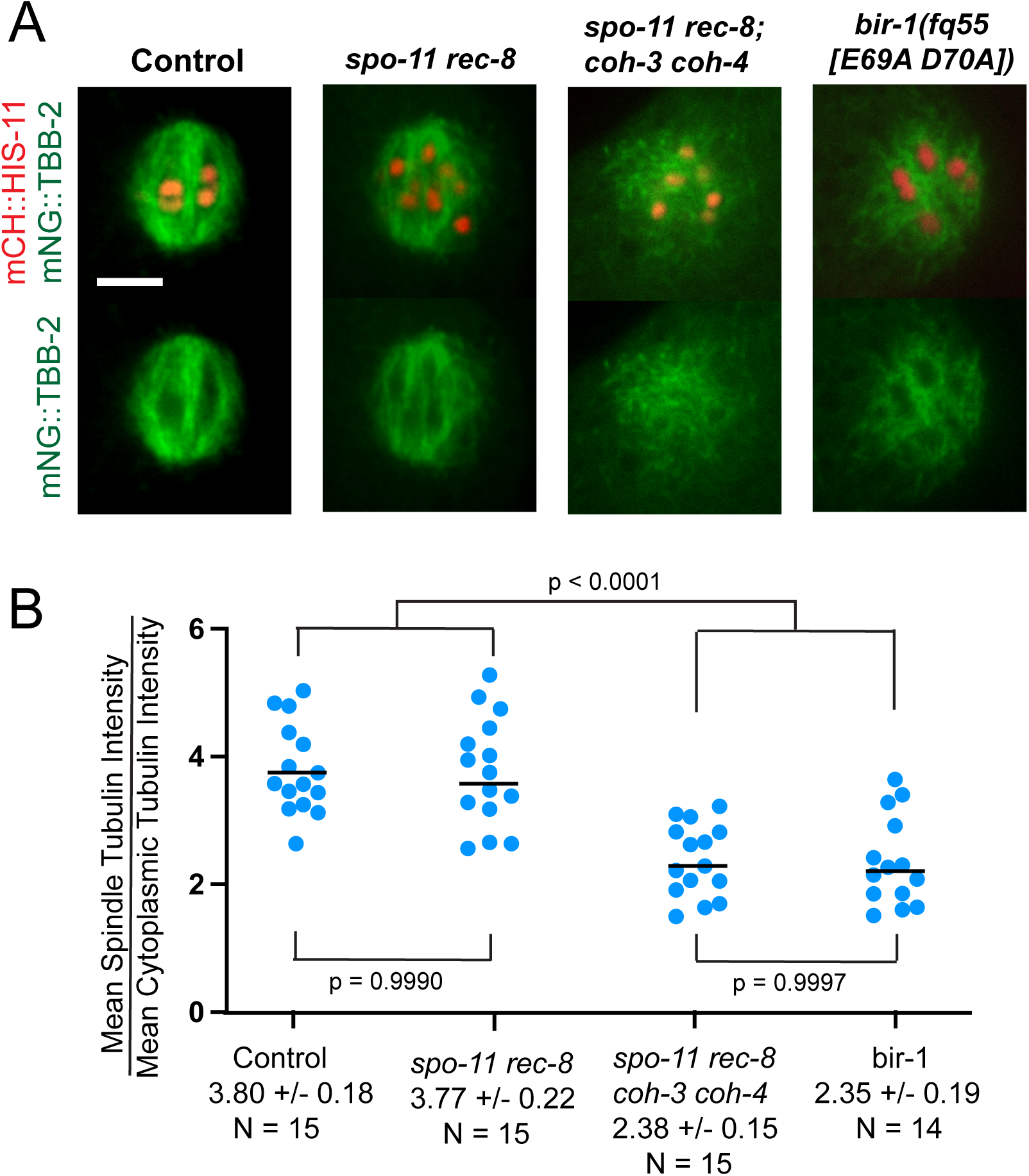
MT density is decreased in *spo-11 rec-8; coh-3 coh-4* and *bir-1(fq55)* spindles. **(A)** Single slices from z-stack images of embryos expressing mNG::TBB-2 and mCH::HIS-11. Bar = 4μm. **(B)** Ratios of mean, background-subtracted mNG::TBB-2 pixel values in spindles vs. nearby cytoplasm of control and mutant embryos. N = number of embryos.

**Supplemental Figure 3.**
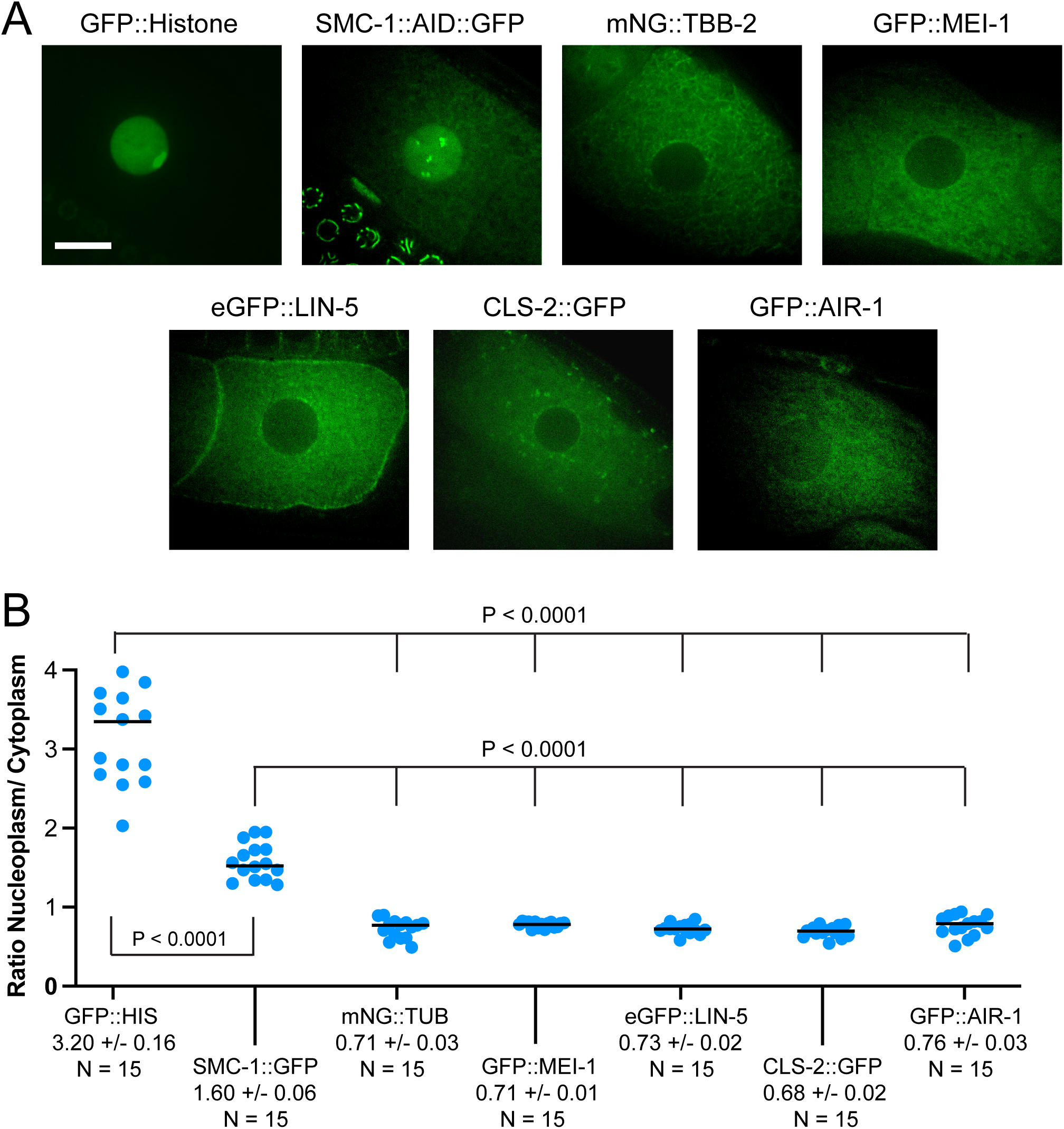
Spindle assembly factors are cytoplasmic prior to nuclear envelope breakdown. **(A)** Single plane images of -1 oocytes in *C. elegans* expressing GFP::H2B, SMC-1::AID::GFP, and spindle assembly factors. Bar = 10 μm. (B) Nucleoplasmic to cytoplasmic ratios were determined for mean, background-subtracted pixel values in -1 oocytes.

## References

Blanco-Ameijeiras, J., Lozano-Fernandez, P., & Marti, E. (2022). Centrosome maturation - in tune with the cell cycle. J Cell Sci, 135(2). doi:10.1242/jcs.259395

Broad, A. J., DeLuca, K. F., & DeLuca, J. G. (2020). Aurora B kinase is recruited to multiple discrete kinetochore and centromere regions in human cells. J Cell Biol, 219(3). doi:10.1083/jcb.201905144

Cahoon, C. K., Helm, J. M., & Libuda, D. E. (2019). Synaptonemal Complex Central Region Proteins Promote Localization of Pro-crossover Factors to Recombination Events During Caenorhabditis elegans Meiosis. Genetics, 213(2), 395–409. doi:10.1534/genetics.119.302625

Carazo-Salas, R. E., Guarguaglini, G., Gruss, O. J., Segref, A., Karsenti, E., & Mattaj, I. W. (1999). Generation of GTP-bound Ran by RCC1 is required for chromatin-induced mitotic spindle formation. Nature, 400(6740), 178–181. doi:10.1038/22133

Castellano-Pozo, M., Pacheco, S., Sioutas, G., Jaso-Tamame, A. L., Dore, M. H., Karimi, M. M., & Martinez-Perez, E. (2020). Surveillance of cohesin-supported chromosome structure controls meiotic progression. Nat Commun, 11(1), 4345. doi:10.1038/s41467-020-18219-9

Cavazza, T., & Vernos, I. (2015). The RanGTP Pathway: From Nucleo-Cytoplasmic Transport to Spindle Assembly and Beyond. Front Cell Dev Biol, 3, 82. doi:10.3389/fcell.2015.00082

Cesario, J., & McKim, K. S. (2011). RanGTP is required for meiotic spindle organization and the initiation of embryonic development in Drosophila. J Cell Sci, 124(Pt 22), 3797–3810. doi:10.1242/jcs.084855

Chuang, C. H., Schlientz, A. J., Yang, J., & Bowerman, B. (2020). Microtubule assembly and pole coalescence: early steps in C aenorhabditis elegans oocyte meiosis I spindle assembly. Biol Open, 9(6). doi:10.1242/bio.052308

Connolly, A. A., Sugioka, K., Chuang, C. H., Lowry, J. B., & Bowerman, B. (2015). KLP-7 acts through the Ndc80 complex to limit pole number in C. elegans oocyte meiotic spindle assembly. J Cell Biol, 210(6), 917–932. doi:10.1083/jcb.201412010

Crawley, O., Barroso, C., Testori, S., Ferrandiz, N., Silva, N., Castellano-Pozo, M., . . . Martinez-Perez, E. (2016). Cohesin-interacting protein WAPL-1 regulates meiotic chromosome structure and cohesion by antagonizing specific cohesin complexes. Elife, 5, e10851. doi:10.7554/eLife.10851

Danlasky, B. M., Panzica, M. T., McNally, K. P., Vargas, E., Bailey, C., Li, W., . . . McNally, F.J. (2020). Evidence for anaphase pulling forces during C. elegans meiosis. J Cell Biol, 219(12). doi:10.1083/jcb.202005179

Deng, M., Suraneni, P., Schultz, R. M., & Li, R. (2007). The Ran GTPase mediates chromatin signaling to control cortical polarity during polar body extrusion in mouse oocytes. Dev Cell, 12(2), 301–308. doi:10.1016/j.devcel.2006.11.008

Divekar, N. S., Davis-Roca, A. C., Zhang, L., Dernburg, A. F., & Wignall, S. M. (2021). A degron-based strategy reveals new insights into Aurora B function in C. elegans. PLoS Genet, 17(5), e1009567. doi:10.1371/journal.pgen.1009567

Dumont, J., Oegema, K., & Desai, A. (2010). A kinetochore-independent mechanism drives anaphase chromosome separation during acentrosomal meiosis. Nat Cell Biol, 12(9), 894–901. doi:10.1038/ncb2093

Dumont, J., Petri, S., Pellegrin, F., Terret, M. E., Bohnsack, M. T., Rassinier, P., . . . Verlhac, M. H. (2007). A centriole- and RanGTP-independent spindle assembly pathway in meiosis I of vertebrate oocytes. J Cell Biol, 176(3), 295–305. doi:10.1083/jcb.200605199

Ellefson, M. L., & McNally, F. J. (2011). CDK-1 inhibits meiotic spindle shortening and dynein- dependent spindle rotation in C. elegans. J Cell Biol, 193(7), 1229–1244. doi:10.1083/jcb.201104008

Ferrandiz, N., Barroso, C., Telecan, O., Shao, N., Kim, H. M., Testori, S., . . . Martinez-Perez, E. (2018). Spatiotemporal regulation of Aurora B recruitment ensures release of cohesion during C. elegans oocyte meiosis. Nat Commun, 9(1), 834. doi:10.1038/s41467-018-03229-5

Gigant, E., Stefanutti, M., Laband, K., Gluszek-Kustusz, A., Edwards, F., Lacroix, B., . . . Dumont, J. (2017). Inhibition of ectopic microtubule assembly by the kinesin-13 KLP-7 prevents chromosome segregation and cytokinesis defects in oocytes. Development, 144(9), 1674–1686. doi:10.1242/dev.147504

Gyuricza, M. R., Manheimer, K. B., Apte, V., Krishnan, B., Joyce, E. F., McKee, B. D., & McKim, K. S. (2016). Dynamic and Stable Cohesins Regulate Synaptonemal Complex Assembly and Chromosome Segregation. Curr Biol, 26(13), 1688–1698. doi:10.1016/j.cub.2016.05.006

Halpin, D., Kalab, P., Wang, J., Weis, K., & Heald, R. (2011). Mitotic spindle assembly around RCC1-coated beads in Xenopus egg extracts. PLoS Biol, 9(12), e1001225. doi:10.1371/journal.pbio.1001225

Heald, R., Tournebize, R., Blank, T., Sandaltzopoulos, R., Becker, P., Hyman, A., & Karsenti, E. (1996). Self-organization of microtubules into bipolar spindles around artificial chromosomes in Xenopus egg extracts. Nature, 382(6590), 420–425. doi:10.1038/382420a0

Jiang, K., Rezabkova, L., Hua, S., Liu, Q., Capitani, G., Altelaar, A. F. M., . . . Akhmanova, A. (2017). Microtubule minus-end regulation at spindle poles by an ASPM-katanin complex. Nat Cell Biol, 19(5), 480–492. doi:10.1038/ncb3511

Joly, N., Beaumale, E., Van Hove, L., Martino, L., & Pintard, L. (2020). Phosphorylation of the microtubule-severing AAA+ enzyme Katanin regulates C. elegans embryo development. J Cell Biol, 219(6). doi:10.1083/jcb.201912037

Kelly, A. E., Ghenoiu, C., Xue, J. Z., Zierhut, C., Kimura, H., & Funabiki, H. (2010). Survivin reads phosphorylated histone H3 threonine 3 to activate the mitotic kinase Aurora B. Science, 330(6001), 235–239. doi:10.1126/science.1189505

Kelly, A. E., Sampath, S. C., Maniar, T. A., Woo, E. M., Chait, B. T., & Funabiki, H. (2007). Chromosomal enrichment and activation of the aurora B pathway are coupled to spatially regulate spindle assembly. Dev Cell, 12(1), 31–43. doi:10.1016/j.devcel.2006.11.001

Kirby, C., Kusch, M., & Kemphues, K. (1990). Mutations in the par genes of Caenorhabditis elegans affect cytoplasmic reorganization during the first cell cycle. Dev Biol, 142(1), 203–215. doi:10.1016/0012-1606(90)90164-e

Laband, K., Le Borgne, R., Edwards, F., Stefanutti, M., Canman, J. C., Verbavatz, J. M., & Dumont, J. (2017). Chromosome segregation occurs by microtubule pushing in oocytes. Nat Commun, 8(1), 1499. doi:10.1038/s41467-017-01539-8

Loughlin, R., Wilbur, J. D., McNally, F. J., Nedelec, F. J., & Heald, R. (2011). Katanin contributes to interspecies spindle length scaling in Xenopus. Cell, 147(6), 1397–1407. doi:10.1016/j.cell.2011.11.014

Mains, P. E., Kemphues, K. J., Sprunger, S. A., Sulston, I. A., & Wood, W. B. (1990). Mutations affecting the meiotic and mitotic divisions of the early Caenorhabditis elegans embryo. Genetics, 126(3), 593–605. doi:10.1093/genetics/126.3.593

Maresca, T. J., Groen, A. C., Gatlin, J. C., Ohi, R., Mitchison, T. J., & Salmon, E. D. (2009). Spindle assembly in the absence of a RanGTP gradient requires localized CPC activity. Curr Biol, 19(14), 1210–1215. doi:10.1016/j.cub.2009.05.061

McCarter, J., Bartlett, B., Dang, T., & Schedl, T. (1999). On the control of oocyte meiotic maturation and ovulation in Caenorhabditis elegans. Dev Biol, 205(1), 111–128. doi:10.1006/dbio.1998.9109

McNally, K., Audhya, A., Oegema, K., & McNally, F. J. (2006). Katanin controls mitotic and meiotic spindle length. J Cell Biol, 175(6), 881–891. doi:10.1083/jcb.200608117

McNally, K. P., & McNally, F. J. (2011). The spindle assembly function of Caenorhabditis elegans katanin does not require microtubule-severing activity. Mol Biol Cell, 22(9), 1550–1560. doi:10.1091/mbc.E10-12-0951

Monen, J., Maddox, P. S., Hyndman, F., Oegema, K., & Desai, A. (2005). Differential role of CENP-A in the segregation of holocentric C. elegans chromosomes during meiosis and mitosis. Nat Cell Biol, 7(12), 1248–1255. doi:10.1038/ncb1331

Mullen, T. J., & Wignall, S. M. (2017). Interplay between microtubule bundling and sorting factors ensures acentriolar spindle stability during C. elegans oocyte meiosis. PLoS Genet, 13(9), e1006986. doi:10.1371/journal.pgen.1006986

Muscat, C. C., Torre-Santiago, K. M., Tran, M. V., Powers, J. A., & Wignall, S. M. (2015). Kinetochore-independent chromosome segregation driven by lateral microtubule bundles. Elife, 4, e06462. doi:10.7554/eLife.06462

Nasmyth, K. (2002). Segregating sister genomes: the molecular biology of chromosome separation. Science, 297(5581), 559–565. doi:10.1126/science.1074757

O’Connell, C. B., Loncarek, J., Hergert, P., Kourtidis, A., Conklin, D. S., & Khodjakov, A. (2008). The spindle assembly checkpoint is satisfied in the absence of interkinetochore tension during mitosis with unreplicated genomes. J Cell Biol, 183(1), 29–36. doi:10.1083/jcb.200801038

Paix, A., Folkmann, A., & Seydoux, G. (2017). Precision genome editing using CRISPR-Cas9 and linear repair templates in C. elegans. Methods, 121-122, 86-93. doi:10.1016/j.ymeth.2017.03.023

Pasierbek, P., Jantsch, M., Melcher, M., Schleiffer, A., Schweizer, D., & Loidl, J. (2001). A Caenorhabditis elegans cohesion protein with functions in meiotic chromosome pairing and disjunction. Genes Dev, 15(11), 1349–1360. doi:10.1101/gad.192701

Radford, S. J., Jang, J. K., & McKim, K. S. (2012). The chromosomal passenger complex is required for meiotic acentrosomal spindle assembly and chromosome biorientation. Genetics, 192(2), 417–429. doi:10.1534/genetics.112.143495

Sampath, S. C., Ohi, R., Leismann, O., Salic, A., Pozniakovski, A., & Funabiki, H. (2004). The chromosomal passenger complex is required for chromatin-induced microtubule stabilization and spindle assembly. Cell, 118(2), 187–202. doi:10.1016/j.cell.2004.06.026

Schlientz, A. J., & Bowerman, B. (2020). C. elegans CLASP/CLS-2 negatively regulates membrane ingression throughout the oocyte cortex and is required for polar body extrusion. PLoS Genet, 16(10), e1008751. doi:10.1371/journal.pgen.1008751

Severson, A. F., Ling, L., van Zuylen, V., & Meyer, B. J. (2009). The axial element protein HTP-3 promotes cohesin loading and meiotic axis assembly in C. elegans to implement the meiotic program of chromosome segregation. Genes Dev, 23(15), 1763–1778. doi:10.1101/gad.1808809

Severson, A. F., & Meyer, B. J. (2014). Divergent kleisin subunits of cohesin specify mechanisms to tether and release meiotic chromosomes. Elife, 3, e03467. doi:10.7554/eLife.03467

Speliotes, E. K., Uren, A., Vaux, D., & Horvitz, H. R. (2000). The survivin-like C. elegans BIR- 1 protein acts with the Aurora-like kinase AIR-2 to affect chromosomes and the spindle midzone. Mol Cell, 6(2), 211–223. doi:10.1016/s1097-2765(00)00023-x

Srayko, M., Buster, D. W., Bazirgan, O. A., McNally, F. J., & Mains, P. E. (2000). MEI-1/MEI- 2 katanin-like microtubule severing activity is required for Caenorhabditis elegans meiosis. Genes Dev, 14(9), 1072–1084. Retrieved from https://www.ncbi.nlm.nih.gov/pubmed/10809666 https://www.ncbi.nlm.nih.gov/pmc/articles/PMC316576/pdf/x2.pdf

Srayko, M., O’toole, E. T., Hyman, A. A., & Müller-Reichert, T. (2006). Katanin disrupts the microtubule lattice and increases polymer number in C. elegans meiosis. Curr Biol, 16(19), 1944–1949. doi:10.1016/j.cub.2006.08.029

Sumiyoshi, E., Fukata, Y., Namai, S., & Sugimoto, A. (2015). Caenorhabditis elegans Aurora A kinase is required for the formation of spindle microtubules in female meiosis. Mol Biol Cell, 26(23), 4187–4196. doi:10.1091/mbc.E15-05-0258

van der Voet, M., Berends, C. W., Perreault, A., Nguyen-Ngoc, T., Gonczy, P., Vidal, M., . . . van den Heuvel, S. (2009). NuMA-related LIN-5, ASPM-1, calmodulin and dynein promote meiotic spindle rotation independently of cortical LIN-5/GPR/Galpha. Nat Cell Biol, 11(3), 269–277. doi:10.1038/ncb1834

Wang, F., Dai, J., Daum, J. R., Niedzialkowska, E., Banerjee, B., Stukenberg, P. T., . . . Higgins, J. M. (2010). Histone H3 Thr-3 phosphorylation by Haspin positions Aurora B at centromeres in mitosis. Science, 330(6001), 231–235. doi:10.1126/science.1189435

Wignall, S. M., & Villeneuve, A. M. (2009). Lateral microtubule bundles promote chromosome alignment during acentrosomal oocyte meiosis. Nat Cell Biol, 11(7), 839–844. doi:10.1038/ncb1891

Willems, E., Dedobbeleer, M., Digregorio, M., Lombard, A., Lumapat, P. N., & Rogister, B. (2018). The functional diversity of Aurora kinases: a comprehensive review. Cell Div, 13, 7. doi:10.1186/s13008-018-0040-6

Woglar, A., Yamaya, K., Roelens, B., Boettiger, A., Kohler, S., & Villeneuve, A. M. (2020). Quantitative cytogenetics reveals molecular stoichiometry and longitudinal organization of meiotic chromosome axes and loops. PLoS Biol, 18(8), e3000817. doi:10.1371/journal.pbio.3000817

Yamagishi, Y., Honda, T., Tanno, Y., & Watanabe, Y. (2010). Two histone marks establish the inner centromere and chromosome bi-orientation. Science, 330(6001), 239–243. doi:10.1126/science.1194498

Yang, H. Y., Mains, P. E., & McNally, F. J. (2005). Kinesin-1 mediates translocation of the meiotic spindle to the oocyte cortex through KCA-1, a novel cargo adapter. J Cell Biol, 169(3), 447–457. doi:10.1083/jcb.200411132

Yang, H. Y., McNally, K., & McNally, F. J. (2003). MEI-1/katanin is required for translocation of the meiosis I spindle to the oocyte cortex in C elegans. Dev Biol, 260(1), 245–259. doi:10.1016/s0012-1606(03)00216-1

Yueh, W. T., Singh, V. P., & Gerton, J. L. (2021). Maternal Smc3 protects the integrity of the zygotic genome through DNA replication and mitosis. Development, 148(24). doi:10.1242/dev.199800

